# Mechanical cues organize planar cell polarity during vertebrate morphogenesis and embryonic wound repair

**DOI:** 10.1101/2025.05.20.654909

**Authors:** Chih-Wen Chu, Satheeja Velayudhan, Jakob H. Schauser, Sapna Krishnakumar, Stephanie Yang, Keiji Itoh, Dominique Alfandari, Ala Trusina, Sergei Y. Sokol

**Affiliations:** Department of Cell, Developmental and Regenerative Biology, Icahn School of Medicine at Mount Sinai, New York, USA

**Author notes:** Department of Veterinary and Animal Sciences, University of Massachusetts Amherst (Dominique Alfandari). Niels Bohr Institute, University of Copenhagen, Denmark (Jakob H. Schauser, Ala Trusina). Equal contribution (Chih-Wen Chu, Satheeja Velayudhan). Correspondence: Sergei Y. Sokol.

**Keywords:** Epithelial morphogenesis, blastopore, neural tube closure, Xenopus embryo, planar cell polarity, mechanical force sensing, actomyosin, ADIP/SSX2IP/Msd1, Diversin/Ankrd6

## Abstract

Planar cell polarity (PCP) is a phenomenon of coordinated cell orientation in many epithelia and is required for early morphogenetic events, such as vertebrate gastrulation or neural tube closure that place embryonic tissues in their proper locations. Known PCP complexes segregate to opposite edges of each cell due to regulatory feedback interactions, however, whether and how PCP is connected to the tension-sensing machinery has been elusive. Here we observed dynamic polarization of Afadin- and α-Actinin-interacting protein (ADIP) in the epithelia adjacent to the involuting marginal zone and the folding neural plate of *Xenopus* embryos, suggesting that it is controlled by mechanical cues. Supporting this hypothesis, ADIP puncta relocated in response to the pulling forces of neighboring ectoderm cells undergoing apical constriction. Moreover, ADIP puncta rapidly polarized in embryos subjected to stretching and during embryonic wound healing. ADIP formed a mechanosensitive complex with the PCP protein Diversin that was distinct from known core PCP complexes and required for wound repair. We propose that mechanically controlled planar polarization of the ADIP-Diversin complex guides cell behaviors in normal morphogenesis and during wound healing.

**Highlights:**

- ADIP is a mechanosensing protein that is planar polarized together with Diversin in epithelia during morphogenesis and wound repair
- Epithelial wound healing redistributes ADIP-containing complexes towards the wound.
- ADIP and Diversin are interdependent and form a complex regulating morphogenesis and wound healing
- Positive regulatory feedback between mechanical forces and PCP signaling drives morphogenesis

## Introduction

Collective tissue movements during embryonic development such as neural tube closure in vertebrates are regulated by mechanical and molecular stimuli. Cells sense mechanical environmental cues through cell-cell and cell-matrix adhesion complexes linked to actomyosin bundles. Forces transmitted through these cytoskeletal networks are converted into biochemical signals that modulate gene expression and protein dynamics ^1,2^. Despite numerous examples of physical forces affecting the localization and signaling of many proteins ^3–5^, the process of mechanotransduction, from force generation to force sensing and transmission between cells, remains poorly understood at the molecular level. Stretching of mechanosensitive proteins, such as F-actin ^6^, α-catenin ^7^, and vinculin ^8,9^, results in the appearance of new protein binding sites. For example, stretched actin filaments recruit actin-binding proteins such as α-Actinin and Myosin II ^10–12^. Actomyosin contractile forces are transmitted from cell to cell via adherens junctions ^13,14^, but how the response to force is amplified and coordinated within the tissue over large distances is largely unknown.

One molecular pathway that may regulate long-range force transmission during morphogenesis is planar cell polarity (PCP) signaling. PCP is a biological *phenomenon* that refers to coordinated polarity of cells in the plane of the tissue, whereas PCP signaling refers to the *molecular mechanisms* underlying planar polarity axis formation and the control of collective cell behaviors. The ‘core PCP’ proteins form signaling complexes segregating to opposite ends of the cell due to regulatory feedbacks, leading to signal amplification ^15–18^. In vertebrate ectoderm derivatives, including the skin and the neural plate, core PCP complexes polarize to the anterior edge of cells ^19–21^ and are genetically required for morphogenesis such as neural tube closure ^22,23^. In addition to core PCP proteins, PCP can be organized by other molecules such as Fat and Dachsous ^18,24,25^, Par3 and Myosin II during germ band extension ^26^, or Fat2 and Lar phosphatase in the fly follicular cell collective migration ^27^.

Consistent with a role of mechanical signals in the establishment of PCP, global patterning signals and tissue deformation have been associated with PCP regulation ^28–33^. Also, the segregation of core PCP complexes in *Ciona* and *Xenopus* depends on Myosin II activity ^20,34^, and the core PCP axis can be altered by ectopic pulling forces ^29,31,35^. While the published work implicates mechanical forces in PCP establishment, the remaining challenge is to understand how PCP signaling senses mechanical forces and converts them into biochemical signals underlying collective cell behaviors.

We previously identified ADIP (Afadin- and α-Actinin-interacting protein), also known as SSX2IP or Msd1 ^36–38^, as physically associated with a mechanosensitive LIM domain protein ^39^. Since ADIP also interacts with two other mechanosensitive proteins, Afadin and α-Actinin ^12,37,40^, we suspected that ADIP may be similarly involved in force sensing. Here we demonstrate that ADIP polarizes in response to tensile forces in an α-Actinin-dependent manner, in the folding neural plate, during wound healing and blastopore closure. Importantly, we show that the core PCP protein Diversin is the interacting partner of ADIP needed to establish tension-induced PCP. Our findings support an essential role of the ADIP-Diversin complex in sensing mechanical cues during wound healing and embryonic morphogenesis.

## Results

To investigate the involvement of ADIP in mechanosensing, we asked how ADIP is distributed in *Xenopus* embryonic ectoderm. In gastrula ectoderm, GFP-tagged ADIP is localized to nonpolarized puncta (**Fig. 1A-1C**, see **Fig. S1** for quantification). Intriguingly, we observed planar polarization of ADIP puncta in the epidermis surrounding the closing neural plate (**Fig. 1D-1G**) and in the cells abutting the blastopore lip during gastrulation (**Fig. 1H-1N**). These locations are adjacent to the areas undergoing apical constriction and likely experience tensile stresses ^22,41,42^. ADIP puncta were predominantly enriched near the cell corners and junctions proximal to the constricting cells, and this polarized distribution was observed in three to six rows of cells adjacent to the constricted cells. Of note, the puncta were consistently polarized towards the neural plate border at different locations along the neural plate (**Fig. 1E-1F’**). Importantly, this ADIP polarity was sensitive to Myosin II inhibition by constitutively active myosin phosphatase Mypt1 ^43,44^(**Fig. S2A-S2D**). We thus hypothesized that ADIP localization is controlled by mechanical forces.

**Fig. 1.**
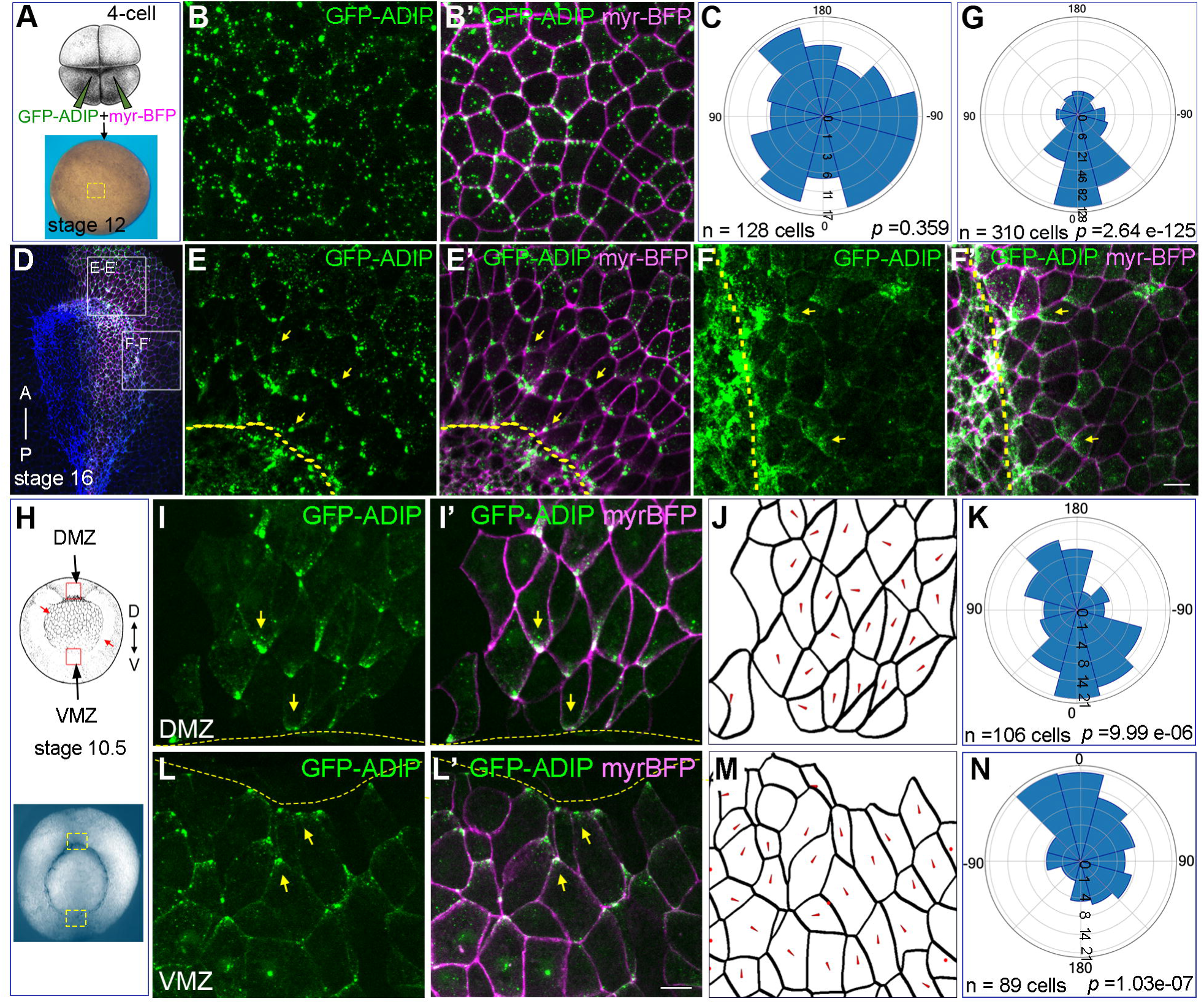
ADIP is polarized in superficial cell layers during morphogenetic events. (A) Scheme of RNA injections. myr-BFP marks the cell boundary. RNAs were targeted to the neural plate and the blastopore of four-cell *Xenopus* embryos as described in the Methods. (B-B’) Localization of GFP-ADIP in ectodermal cells at the boxed region in (A) at stage 12. (C) Rose plot showing ADIP orientation in ectoderm. Chi-square test indicates random distribution. Also see Figure S1. (D-F’) Top view of stage 16 anterior neural plate expressing GFP-ADIP and myr-BFP. The neural plate is highlighted by intense staining of F-actin (blue). Anterior (E, E’) and lateral (F, F’) cells at the respective boxed regions in (D) show ADIP enrichment at the cell corners (arrows) closest to the neural plate border (dashed lines). Also see Figure S2. (G) Rose plot showing ADIP orientation with respect to the anterior neural plate border (0°). Chi-square test indicates non-random ADIP distribution (*p* <0.05). (H) Vegetal view of *Xenopus* gastrula at stage 10.5 depicting the blastopore lips (red arrows). (I, I’, L, L’) Superficial cells from the boxed regions in (H) show ADIP enrichment (arrows) toward the blastopore lip boundary (dashed lines). Scale bars, 20 μm. (J, M) Segmentation of cells in (I, I’) and (L, L’), respectively, showing ADIP polarization (red arrowheads). (K, N) Rose plots showing ADIP orientation with respect to the blastopore lip (0°) in (I, I’) and (L, L’) respectively. Data are from 3 embryos. Chi-square test indicates non-random ADIP distribution.

To evaluate force-sensing properties of ADIP, we asked whether it can relocalize in response to external mechanical cues. To produce a local pulling force, ectodermal cells were injected with mRNA encoding Shroom3, an apical constriction inducer ^45,46^. While GFP-ADIP puncta were randomly distributed in control embryonic ectoderm (**Fig. 2A-2C**), they accumulated near the cell borders adjacent to the mosaic patches of neighboring Shroom3-expressing cells (**Fig. 2D-2F**). Live imaging revealed the appearance of ADIP puncta near the apical surface and their movement toward these cell borders (**Fig. S3A** and **Video S1**), suggesting directional transport of ADIP puncta. Similar results have been obtained in experiments using Plekhg5, another apical constriction inducer ^47^ (**Fig. S3B-S3G**), indicating that this response is not specific to Shroom3 but rather is triggered by apical constriction *per se*.

**Fig. 2.**
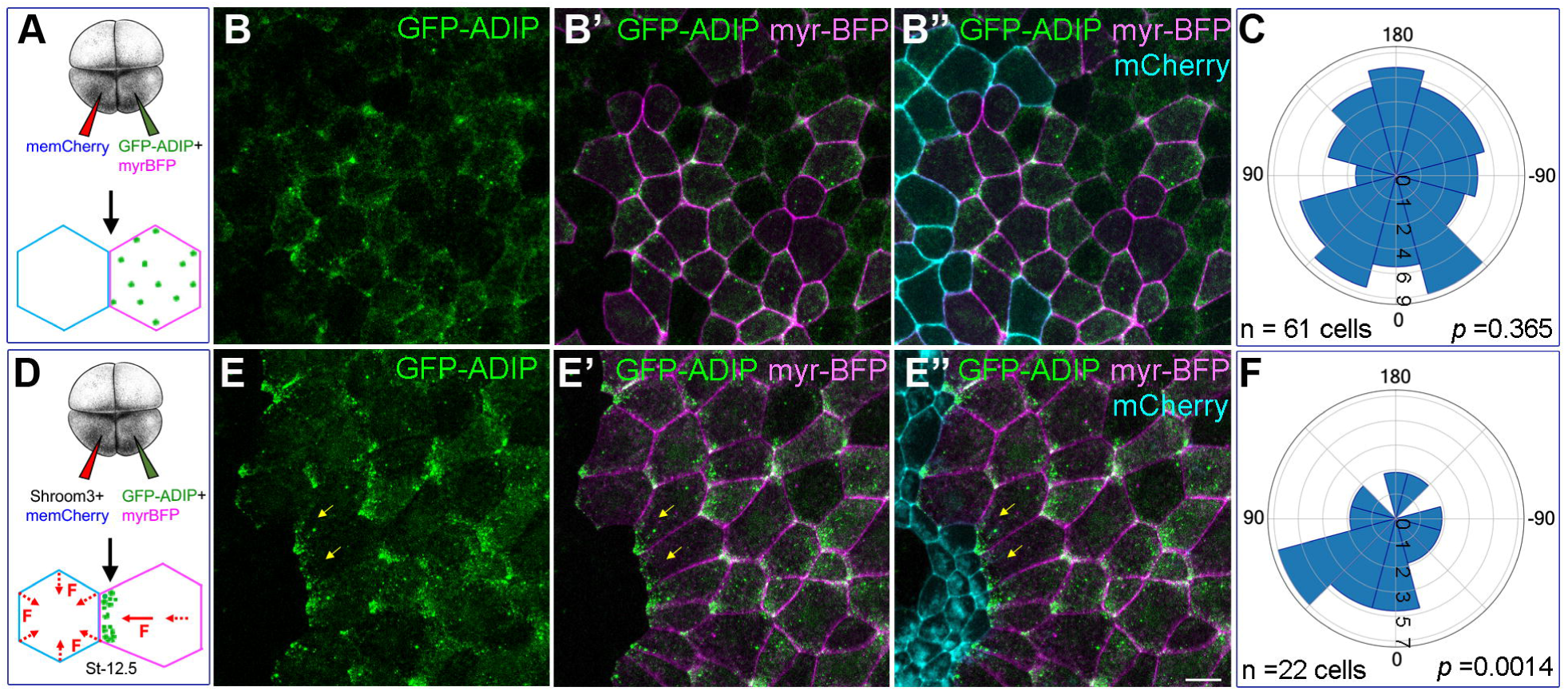
ADIP polarization in response to anisotropic mechanical forces. (A, D) Scheme of injection. The clone of cells injected with Shroom3 RNA or control are marked with mem-Cherry, the clone of cells expressing GFP-ADIP is marked by myr-BFP (B-B’’, E-E’’) Distribution of GFP-ADIP (green) in cells adjacent to mem-Cherry-expressing cells (cyan) at stage 12.5. Note the polarization of GFP-ADIP (arrows in E-E”) toward the adjacent mem-Cherry cells expressing Shroom3 (cyan). (C, F) Rose plots showing ADIP orientation with respect to adjacent mem-Cherry-expressing cells (0°) in (B-B”) and (E-E”) respectively. Data are from 3 embryos. Chi-square test indicates random (B-B”) and non-random (E-E”) distribution, respectively. Bar, 20 µm. Also see Figure S3.

To further confirm that ADIP can polarize in response to tensile forces along the tissue plane, embryos expressing GFP-ADIP were subjected to unidirectional stretching along the animal-vegetal axis using a hydrostatic pressure-driven aspiration device ^29^ (**Fig. 3A**). ADIP puncta were randomly distributed in the control unmanipulated embryos (**Fig. 3B-3D**) but became polarized along the force vector in response to a 25-min stretch (**Fig. 3E-3G**). Like ADIP polarization around the neural plate, the stretch-induced puncta polarization was sensitive to Mypt1 (**Fig. S4A-S4H**), consistent with the requirement of Myosin II for ADIP polarization. Together, our observations strongly indicate that ADIP localization reflects the existing mechanical cues along the tissue plane.

**Fig. 3.**
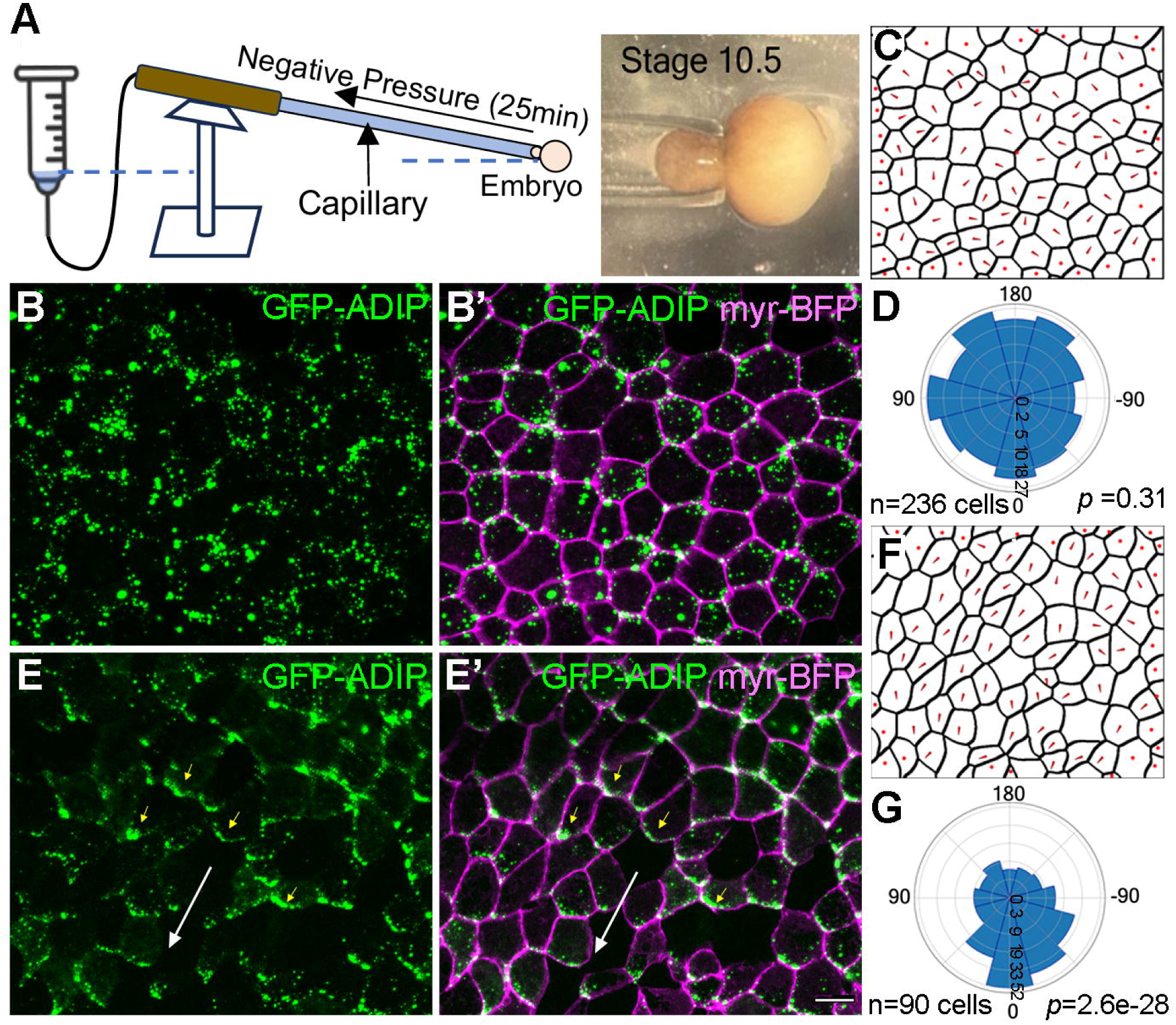
ADIP redistribution in response to aspiration-induced stretching. (A) Experimental setup of unidirectional tissue stretching. Negative pressure was applied to gastrula embryos to aspirate ectoderm into the capillary. (B-B’, E-E’) GFP-ADIP distribution in an unstretched (B-B’) or stretched embryo (E-E’). Note the polarization of GFP-ADIP (yellow arrows) that aligns with the stretch axis (white arrow) (E-E’). Scale bar, 20 μm. (C, F) Segmentation of cells in (B-B’) and (E-E’) respectively showing ADIP polarity (red arrowheads). (D, G) Rose plots showing ADIP orientation with respect to the animal pole or aspiration axis (0°) in (B-B’) and (E-E’) respectively. Data are from 3 embryos. Chi-square test indicates random distribution in (B-B’) and non-random distribution in (E-E’) respectively. Also see Figure S4.

An alternative model that is easily amenable to experimental manipulation is embryonic wound healing that depends on the pulling force of the contractile actomyosin ring surrounding the wound ^48–52^. Consistent with our previous conclusions, we observed robust polarization of ADIP puncta within 20-40 min after inflicting a wound by surgically removing a piece of gastrula ectoderm at stage 11 (**Fig. 4A-4F, Fig. S5A-S5B”’,** and **Video S2).** Although the wound repair can involve both superficial and deep cell layers of *Xenopus* ectoderm ^52,53^, we observed ADIP polarization even in case of exclusively superficial wounds. Importantly, ADIP was polarized in 5-9 rows of cells surrounding the closing wound, with the degree of polarization (angular alignment, i.e. ‘orientation’, and magnitude) decreased in the cells that are further away from the wound edge (**Fig. S5C-S5F**). Moreover, time-lapse imaging revealed directional movements of ADIP puncta (**Fig. 4G, Video S2, arrows**) or appearance of ADIP puncta near the cell corners proximal to the wound (**Video S2, asterisks**). We observed gradual increase of ADIP puncta at the cell corners, whereas the numbers of apical cortical puncta and those near the bicellular junctions gradually decreased (**Fig. 4H**). These data suggest that ADIP polarization is likely achieved via directional transport of ADIP puncta toward the cell corner closest to the wound.

**Fig. 4.**
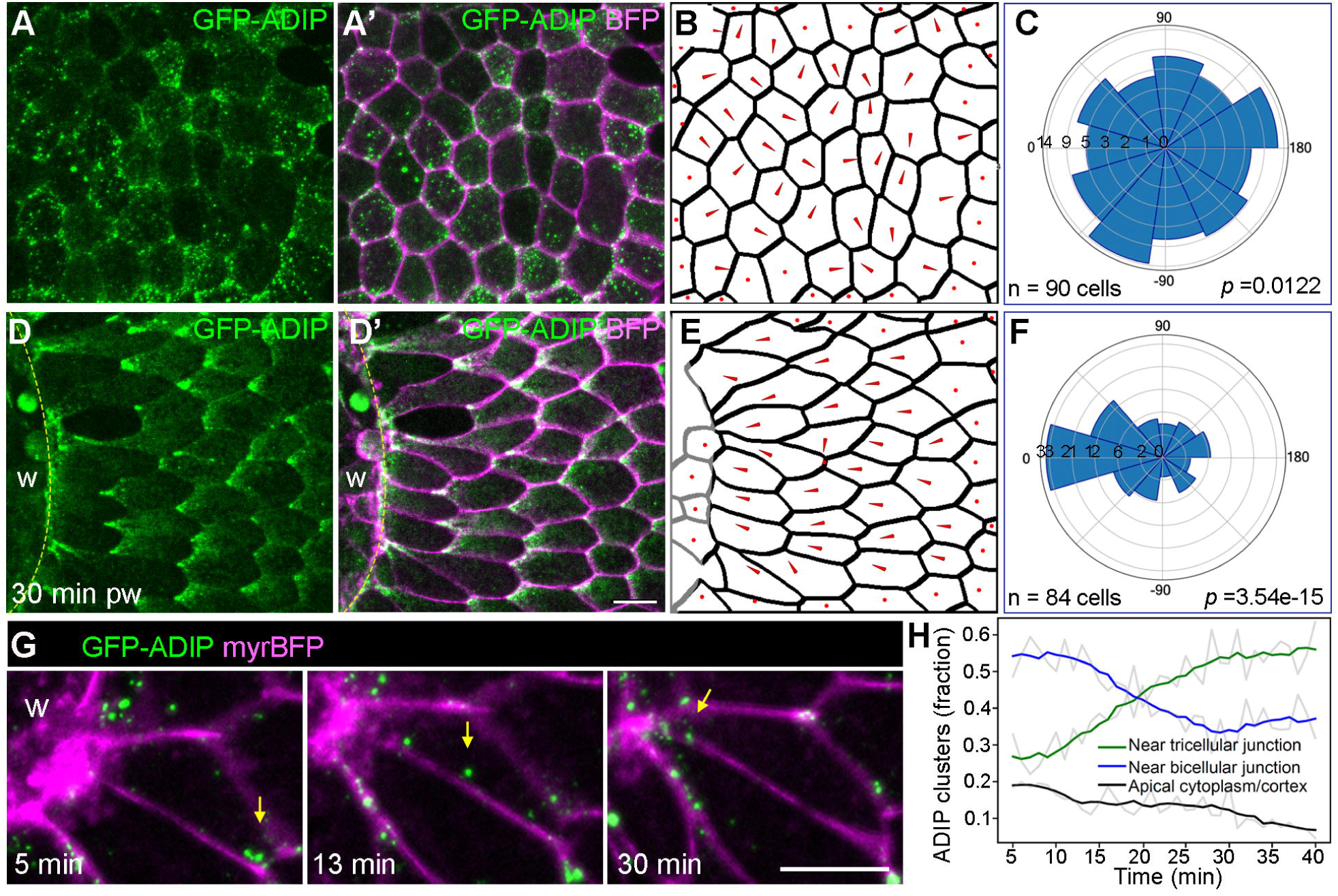
ADIP is polarized in epithelial cells around the wound. (A-A’, D-D’) Localization of GFP-ADIP in unwounded (A-A’) and 30 min post-wounded (D-D’) gastrula ectoderm respectively, with myr-BFP marking the cell boundaries. Wound (w) edge is indicated by dashed lines. Also see Figure S5A. (B-B’, E-E’) Segmentation of cells in (A-A’) and (D-D’) respectively showing ADIP polarity (red arrowheads). (C, F) Rose plots showing ADIP orientation with respect to the animal pole or wound edge (0°) in (A-A’) and (D-D’) respectively. Data are from 3 embryos. Chi-square test indicates random distribution in (A-A’) and non-random distribution in (D-D’) respectively. Also see Figure S5B-F, S6, S7. (G) Stills of time-lapse images showing GFP-ADIP at 5, 13 and 30 min post-wounding (pw) with myr-BFP marking the cell boundaries. Note ADIP movement toward the wound (w) over time (arrows). Also see Video S2. (H) Quantification of the distribution of ADIP puncta near the bicellular junction, tricellular junction, or apical cytoplasm over time. Colored lines are the rolling averages of the original data points shown in grey.

To test which region of ADIP is required for its polarization, ADIP deletion mutants lacking the amino acid sequences corresponding to the α-Actinin-or Afadin-binding region ^37^ (**Fig. S6A**) were expressed in the wounded ectoderm. We found that the mutant lacking Afadin-binding site remained partly polarized, whereas the mutant lacking the α-Actinin-binding domain failed to polarize toward the wound (**Fig. S6B-S6G**). The latter finding indicates that α-Actinin is involved in ADIP polarization. We next assessed the distribution of F-actin and the Myosin II heavy chain that can be traced by the SF9 intrabody ^54–56^ and detected their polarization towards the wound edge (**Fig. S7A-S7D)**. These findings suggest that ADIP polarization is associated with actomyosin contractility near the cell vertices.

To understand the significance of ADIP in wound healing, we assessed the effect of ADIP knockdown on wound closure. We generated ADIP-specific monoclonal antibodies and confirmed the depletion of endogenous ADIP (65 kDa) in embryos injected with morpholino (MO) targeting ADIP (ADIPMO) ^39^ (**Fig. 5A**). Whereas the epithelial wounds were nearly closed within 15 min in control embryos, the rate of wound closure in ADIP morphants was significantly reduced, especially in the first 5 min after wounding (**Fig. 5B and 5C**). This effect can be rescued by co-expressing GFP-ADIP RNA (**Fig. 5D**), confirming specificity. In search for molecular markers affected by ADIP depletion, we found that F-actin staining was significantly reduced both at bicellular junctions and at the wound edge in ADIP depleted cells as compared to control cells (**Fig. 5E-5G, Fig.S8A-S8C)**. Moreover, junctional E-cadherin was decreased in the depleted tissue (**Fig. S8D-S8H**), suggesting that ADIP acts in the assembly or maintenance of adherens junctions that play a critical role during wound closure ^50^.

**Fig. 5.**
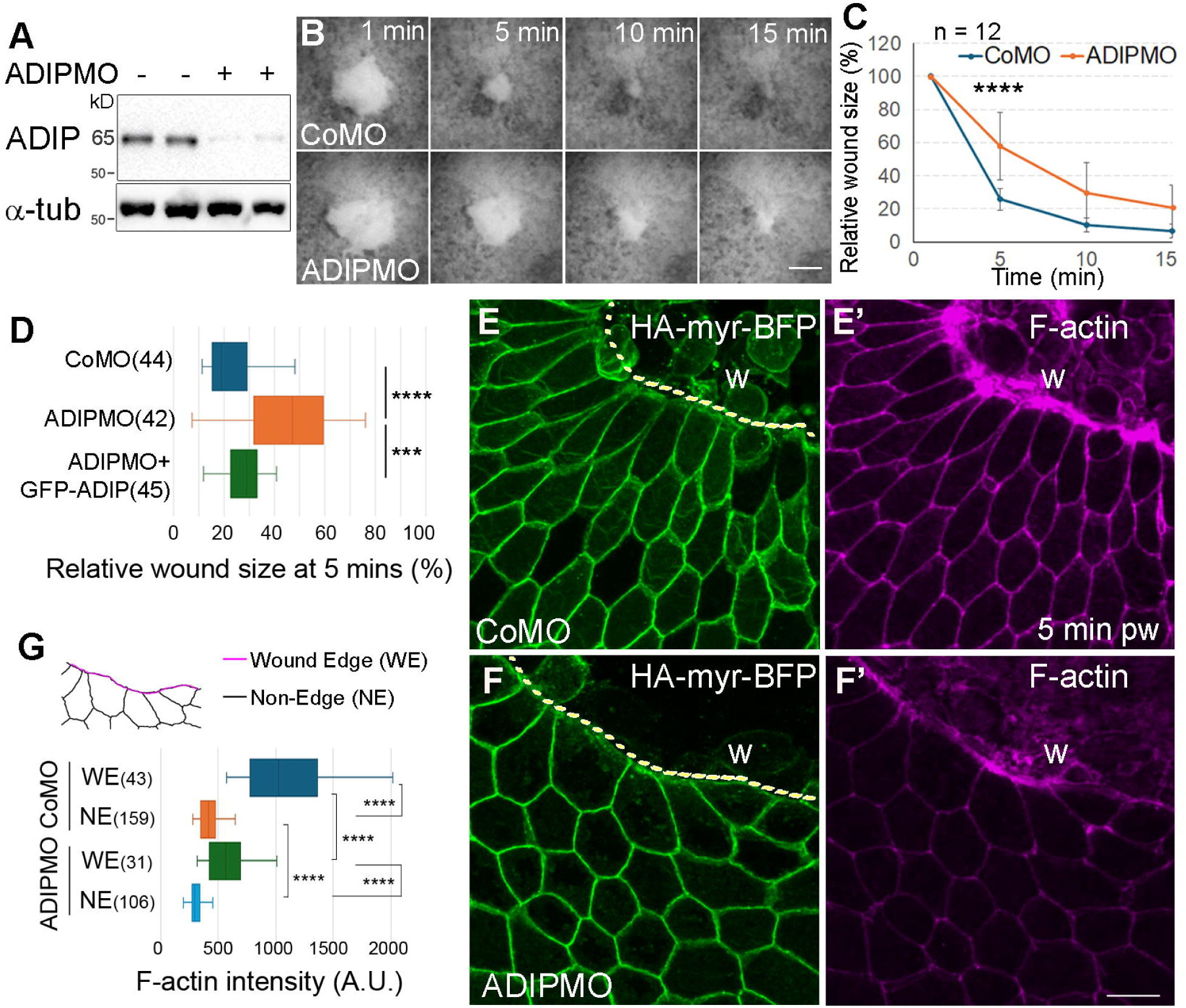
ADIP is required for epithelial wound healing. (A) Immunoblot showing the depletion of endogenous ADIP (65 kDa band) in ADIPMO-injected embryo lysates. α-tubulin (α-tub) is the loading control. (B-D) ADIP depletion delayed wound healing in stage 14 ectoderm. (B) Stills of time-lapse images showing the epithelial wound in CoMO or ADIPMO-injected stage 14 embryos at indicated time point after wounding. (C) Quantification of the wound size relative to that at 1 min post-wounding (relative wound size) over time. Data are from two clutches of embryos. Two-tailed Student’s *t*-test of the relative wound size at 5 mins: ****: *p* <0.0001. (D) Quantification of the relative wound size at 5 min post-wounding showing rescue of the ADIPMO defect by GFP-ADIP. Data are from three clutches of embryos, and *p* values are obtained using one-way ANOVA with Tukey post hoc test. ****: *p* <0.0001, ***: *p* <0.001. (E-H) ADIP depletion reduced F-actin enrichment at the wound edge. Also see Figure S8. (E-F’) Epithelial wounds (w) 5 min post-wounding on embryos injected with indicated MOs were immunostained to visualize HA (green) and F-actin (magenta). HA-myr-BFP is the lineage tracer marking the cell boundaries, dashed lines mark the wound edge. (G) Segmentation of cells and quantification of F-actin intensity at the wound edge (WE, magenta) in (F-F’) and at non-wound edge junctions (NE, black). Data are from three embryos, 5 min post-wounding. Total cell numbers are in parentheses. Two-tailed Student’s t test: ****: *p* <0.0001.

The planar polarized distribution of ADIP is very similar to that of core PCP complexes in many epithelia, raising the question of whether ADIP functions in PCP signaling. We therefore assessed the effects of ADIP depletion on the polarization of the core PCP protein Vangl2 in the neural plate (**Fig. S9A**). In control embryos, Vangl2 accumulated at the anterior edges of neuroepithelial cells at stage 15 as reported previously ^20^(**Fig. S9B and S9B’**, arrows). This enrichment was lost in embryos depleted of ADIP (**Fig. S9C and S9C’,** asterisks). Immunoblotting confirmed that Vangl2 protein level remained unaffected (**Fig. S9D**), suggesting that ADIP regulates Vangl2 protein distribution. We next hypothesized that the distribution of Vangl2 may be altered in the wounded gastrula ectoderm, but no detectable Vangl2 polarization was apparent **(Fig. S9E and S9E’)**.

While our results indicate that ADIP functionally interacts with Vangl2, lack of Vangl2 polarization during wound healing prompted us to look for alternative markers that become polarized near the wound edge. We assessed the distribution of Diversin/Ankrd6, a homolog of *Drosophila* PCP protein Diego ^57,58^, because it was previously reported to polarize in the *Xenopus* neural plate ^59^. Like ADIP, Diversin was enriched at the cell corners proximal to the wound edge in gastrula ectoderm (**Fig. 6A-6C**).

**Fig. 6.**
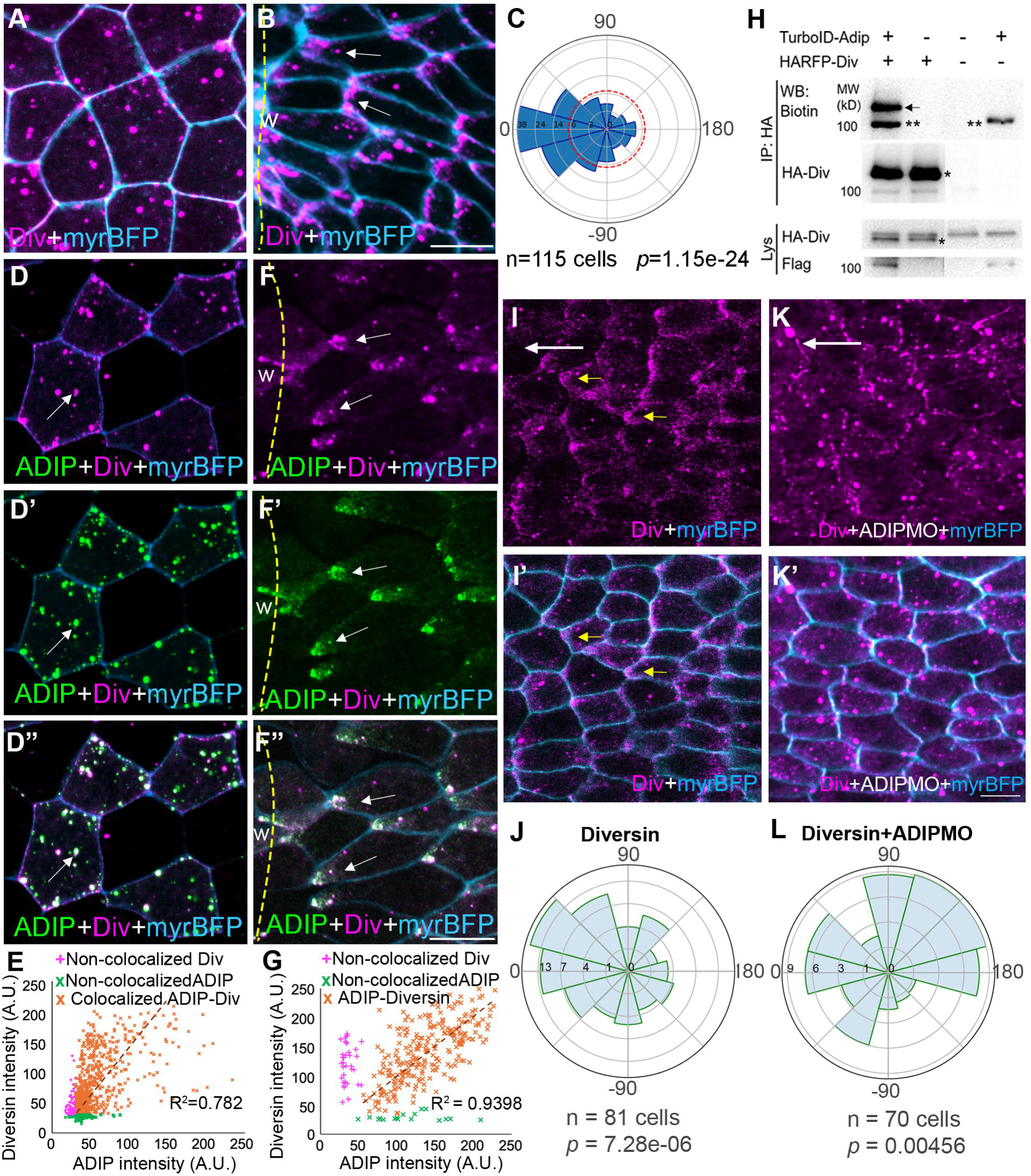
ADIP interacts with Diversin to establish PCP during wound healing. (A-B) Localization of Diversin in stage 11 ectoderm unwounded (A) or 30 min post-wounding (B). Arrows indicate enrichment of Diversin at junctions facing the wound (w) edge (dashed line). (C) Rose plot showing the orientation of Diversin relative to the wound (0°). Data are from 3 embryos. Chi-square test indicates non-random distribution compared to the null hypothesis (dashed line). (D-H) ADIP was associated with Diversin. (D-D”, F-F”) Colocalization of ADIP with Diversin in stage 11 ectoderm, unwounded (D-D”) or 30 min post-wounding (F-F”). Arrows indicate puncta where ADIP and Diversin signals overlap. Dashed lines mark the wound (w) edge. (E, G) Scatter plots showing the colocalization of ADIP and Diversin puncta in (D-D”) and (F-F”) images, respectively. Colocalization of ADIP and Diversin puncta was analyzed using the ComDet plugin in ImageJ and the scatter plots were generated using Microsoft Excel. (E) 74.66% of Diversin puncta colocalized with ADIP, while 15.86% of ADIP puncta and 9.46% of Diversin puncta did not colocalize. (G) At the cell corners facing the wound, 83.12% of Diversin puncta colocalized with ADIP with a strong correlation (*R²*□=□0.9398). (H) Western blot showing immunoprecipitation of HARFP-Diversin (HARFP-Div, asterisks) from embryonic lysates co-expressing TurboID-ADIP and HARFP-Div. Diversin is specifically biotinylated in the presence of TurboID-ADIP (arrow). Biotinylated TurboID-ADIP is also detected in the precipitates (double asterisks). (I-L) ADIP is required for Diversin polarization in response to aspiration-induced stretching. (I-I’, K-K’) RFP-Diversin distribution in stretched embryo with (K-K’) or without (I-I’) ADIPMO (40 ng) injection. Note the polarization of Diversin (yellow arrows) that aligns with the stretch axis (white arrow) in control embryos (I-I’). Scale bar = 20 μm. (J, L) Rescaled rose plots showing the orientation of Diversin polarization relative to the stretch axis (0°) in (I-I’) and (K-K’) respectively. Data are from 3 embryos. Welch’s t-test between population polarities in J and L: *p*<0.0001.

Given the similar distribution of ADIP and Diversin in the epithelium surrounding the wound, we asked whether the two proteins form a complex. Supporting this possibility, the ADIP and DIversin puncta colocalized in the intact gastrula ectoderm (**Fig. 6D-6E**), and the protein complex polarized towards the epidermal wound (**Fig. 6F-6G**). The interaction between ADIP and Diversin was further supported in a proximity-labeling assay, as Diversin was specifically biotinylated by the ADIP construct fused with TurboID ^60^ (**Fig. 6H**). Importantly, the Diversin deletion mutant lacking the ankyrin repeats (Div-ΔANK) ^61^ did not colocalize with ADIP (**Fig. S10A-S10D**) and failed to polarize in response to wounding (**Fig. S10E-S10H).** This finding suggests that Diversin responds to mechanical cues by associating with ADIP. This is further supported by the lack of Diversin polarization in the ectoderm of ADIP-depleted embryos after unidirectional stretching (**Fig. 6I-6L**).

We next asked whether Diversin is required for ADIP polarization because both proteins were implicated in neural tube closure ^39,59^. We also found that wound healing was delayed in embryos depleted Diversin with previously characterized MO ^62^ (**Fig. 7A-7B**). Compared to control embryos, ADIP became less polarized in Diversin-deficient embryos subjected to unidirectional stretching (**Fig. 7C-7F**), indicating that Diversin is required for ADIP polarization. These observations are consistent with ADIP acting together with Diversin in a pathway that is critical for both wound healing and normal morphogenesis.

**Fig. 7.**
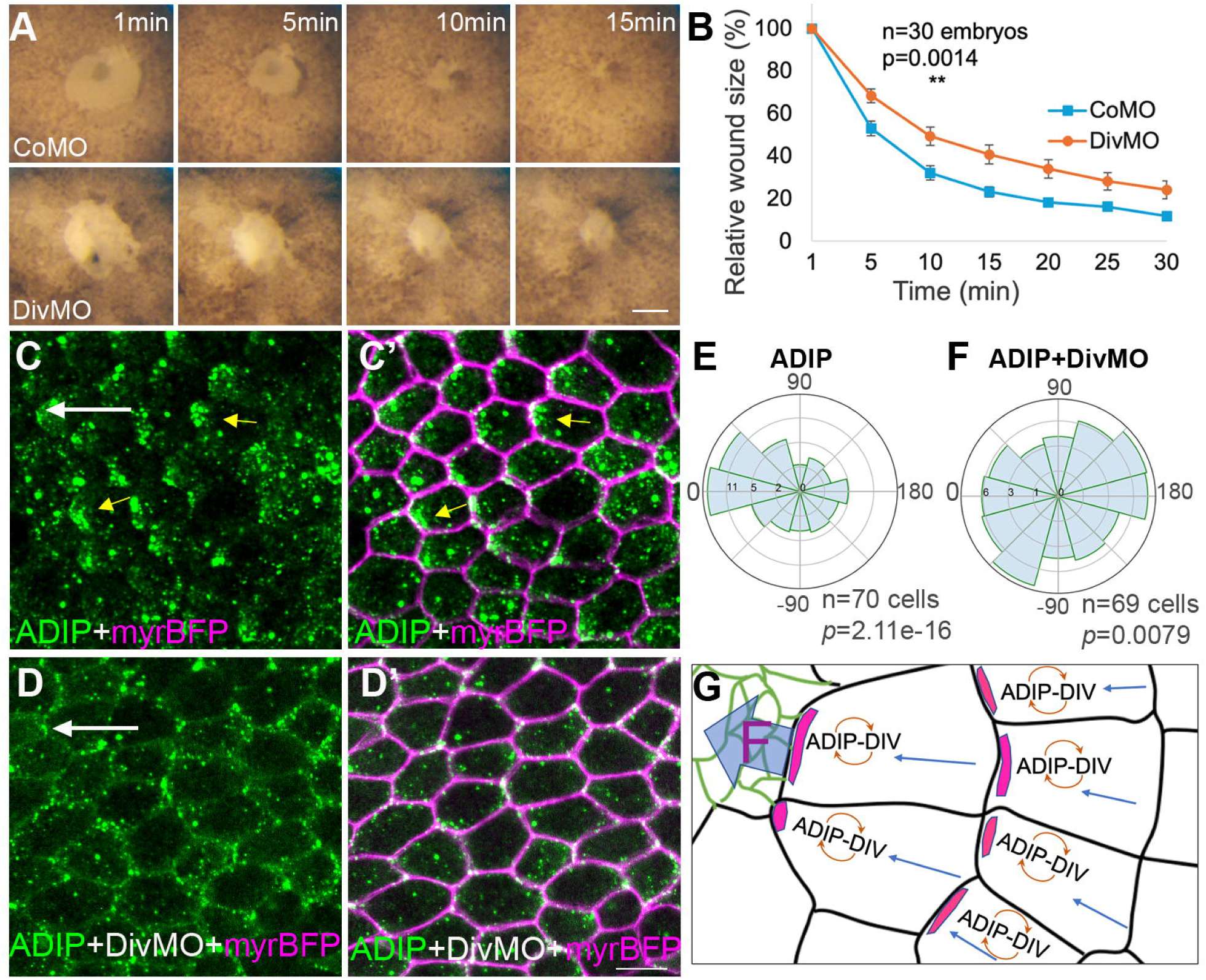
Diversin is required for wound healing and ADIP polarization. (A-B) Diversin requirement for wound healing in stage 11.5 ectoderm. (A) Stills from time-lapse imaging showing epithelial wound closure in CoMO and Diversin MO (DivMO, 60 ng)-injected embryos at the indicated time points post-wounding. Bar, 50 µm. (B) Quantification of the relative wound size over time. Data are from three clutches of embryos. Two-tailed Student’s *t*-test of the relative wound size at 1 min: **: *p* = 0.0014. (C-F) Diversin is required for ADIP polarization in response to aspiration-induced stretching. (C-D′) GFP-ADIP distribution in stretched embryo with (D-D’) or without (C-C’) DivMO (40 ng) injection. Note the polarization of ADIP (yellow arrows) that aligns with the stretch axis (white arrow) in control embryos (C-C’). Scale bar = 20 μm. (E-F) Rescaled rose plots showing the orientation of ADIP polarization relative to the aspiration axis (0°) in (C-C’) and (D-D’) respectively. Data are from 3 embryos. Welch’s t-test between population polarities in E and F: *p*<0.0001. (G) Working model. The ADIP-Diversin (Div) complex (magenta) is redistributed to the proximal end of the cell (blue arrows) in response to the pulling force (F) from neighboring cells. This relocalization leads to the establishment of PCP and subsequent cell-cell signaling in the tissue plane.

## Discussion

In this study we report that ADIP redistributes in response to tensile forces in *Xenopus* embryonic epithelia, leading to the formation of the polarized complex with the PCP protein Diversin. Since both ADIP and Diversin function in morphogenesis and wound healing in *Xenopus* embryos, the enrichment of their complex to one side of the cell may trigger a unique pathway instructing collective cell behaviors yet different from the existing PCP signaling pathways (**Fig. 7G)**. We propose that this protein complex mediates the transmission of tensile forces to biochemical signaling that underlies many morphogenetic processes including gastrulation and neurulation.

During epithelial morphogenesis, cells sense mechanical forces at cell vertices, tension hotspots where the balance between cell adhesion and cortical contractility is fine-tuned to facilitate cell shape changes while maintaining tissue integrity ^63–65^. Here we demonstrate that ADIP redistributes to the proximity of cell vertices in response to tensile force in a Myosin II-dependent manner, and the underlying mechanism may be related to the mechanosensitive properties of ADIP binding partners. Our finding that ADIP polarization requires its α-Actinin-binding domain suggests that ADIP is recruited to the cortex by α-Actinin, a mechanosensitive protein that responds to F-actin deformation ^12^. Moreover, we observe that deletion of the Afadin-binding domain partially affects ADIP polarization, consistent with the mechanosensitive role of Afadin at tricellular junctions ^40,66^. As we did not detect Afadin polarization during wound healing, it remains unclear how Afadin regulates ADIP distribution.

PCP is a hallmark of many epithelial tissues undergoing morphogenesis, and wound healing has significant similarity to morphogenetic processes ^48,67^. Moreover, mouse embryonic skin repair is delayed in the absence of core PCP protein function ^68^, although, to our knowledge, PCP has not been previously described in the wounded epithelium. The localization of the ADIP-Diversin complex appears to be controlled differently from the previously described anterior Vangl2-Prickle complex ^20^. Consistent with this view, the dynamic polarization of Diversin and ADIP takes place in response to wounding at the time when the Vangl2 polarity in the embryo is not yet established ^20,29^. The ADIP-Diversin complex polarization occurs via a unique mechanism that is rapidly activated in response to force. This property makes the ADIP-Diversin polarity module distinct from other polarity modules. We propose that the ADIP-Diversin complex defines a novel mechanosensitive planar polarity system that plays a critical role in wound healing and morphogenesis, adding to the existing PCP signaling modes ^18,24,69^.

To explain the ability of ADIP puncta to move long distances in the cells, we speculate that an additional mechanosensitive trafficking mechanism delivers ADIP in the proximity of the cell vertices. Such directional transport could take place along F-actin filaments or microtubules. At present, there is no evidence for F-actin involvement in ADIP transport. Supporting the potential involvement of microtubules, ADIP mediates microtubule anchorage and can be delivered by microtubule motors Dynein and Kinesin-14 ^36,70,71^. Moreover, extrinsic mechanical cues have been shown to promote microtubule alignment along the force axis ^29^. A hypothesis consistent with the current findings is that the initial recruitment of ADIP by α-Actinin promotes local microtubule anchoring, leading to further accumulation of ADIP through active transport along the microtubules. While the polarized ADIP-Diversin puncta are likely non-centrosomal due to their high number, the association of ADIP and Diversin with the centrosome ^36,61,70,72,73^ may be relevant for their signaling function. For example, the ADIP complex may anchor noncentrosomal microtubules at the polarized cortex to promote PCP-related vesicular trafficking ^59,74,75^.

Our finding of F-actin reduction in ADIP morphants suggests that ADIP not only acts as a force sensor but contributes to subsequent force generation by modulating junctional F-actin assembly. Supporting this hypothesis, we observed polarized distribution of F-actin and Myosin heavy chain toward the closing wound similar to the polarization of ADIP. Future studies will evaluate whether ADIP regulates F-actin through its binding partners Afadin and α-Actinin or through stabilizing junctional E-cadherin. Our data thus suggest a positive feedback loop between mechanical forces produced during morphogenesis and the establishment of polarized ADIP-Diversin signaling complexes that might lead to new rounds of force generation.

## Resource availability

### Lead contact

Requests for further information, resources, and reagents should be directed to and will be fulfilled by the lead contact, Sergei Y. Sokol.

### Materials availability

Reagents generated for this study are available upon reasonable request from the lead contact.

### Data and code availability

All of the original data and any additional information required to reanalyze the data reported in this paper are available from the lead contact upon request.

## Acknowledgments

We thank Chenbei Chang, Naoto Ueno, Ann Miller, Ken Takemaru for plasmids; Ken Irvine and Vladimir Gelfand for comments on the manuscript, Raymond Keller and Lance Davidson for suggestions regarding the aspiration assay. We are grateful to members of the Sokol laboratory for valuable discussions. This research was supported by the NIH grants R35GM122492 to S.Y.S., R24OD021485 to D.A. and the NNF grant NNF23OC0086722 to A.T.

## Author contributions

C.C. and S.V. designed, executed, and analyzed experiments, prepared the figures, and contributed to writing of the manuscript. J.H.S. and A.T. performed image analysis and quantification, statistical analysis, and contributed to writing of the manuscript. S.K. carried out the experiments with stretching embryos and their imaging. S.Y. contributed to wound healing studies of Diversin. D.A. generated ADIP-specific hybridomas and selected monoclonal antibodies to ADIP. K.I. prepared ADIP antigens for immunization and tested the antibody to ADIP by immunoblotting. S.Y.S. conceived the study, designed experiments, acquired funding, supervised and coordinated the project, and wrote the manuscript. All authors agreed to the final version of the manuscript.

## Declaration of generative AI and AI-assisted technologies in the writing process

During the preparation of this work, the authors did not use AI-assisted technologies.

## Supplemental Information

**Document S1**. Figures S1–S10.

**Video S1.** ADIP polarization in response to apical constriction of neighboring cells, related to Figure 2.

Time-lapse images of ectodermal cells at stage 12 showing directional movement of RFP-ADIP (red) toward the apically constricting Shroom3-expressing cells (cyan). Images were taken every 2 mins.

**Video S2.** ADIP polarization in response to wounding, related to Figure 3.

Time-lapse images of gastrula ectoderm at stage 11 showing directional movement of GFP-ADIP (arrows) toward the wound (w). Asterisks mark ADIP puncta appearance near the tricellular junctions. Images were taken every 1 min.

## Methods

### Plasmid constructs

The plasmids encoding pCS105-HA-RFP-Diversin, pCS105-HA-RFP-Diversin-ΔAnk ^58,61^, Mypt1-T696A (the constitutively active mutant of myosin phosphatase) ^43^, pCS2-Plekhg5, a gift of Chenbei Chang ^47^, pCS2-SF9-mNG ^54^, pCS2-myc-mShroom3, pCS2-myr-tagBFP-HA and pCS2-memCherry have been previously described ^59,76^. The *Xenopus* ADIP constructs, pXT7-EGFP-ADIP and pCS105-HA-RFP-ADIP were also described ^39^. FlagTurboID-ADIP has been generated by subcloning of ADIP cDNA insert into the FlagTurboID plasmid (a gift of Kenichi Takemaru, Addgene plasmid 124646)^60^. GFP-Afadin was constructed by subcloning *Xenopus* Afadin cDNA, a gift of Naoto Ueno ^77^, in-frame with GFP into pCS2.

Xenopus ADIP deletion mutants ADIP-ΔAB (actinin-binding domain, aa 124-214) and ADIP-ΔAFB (afadin-binding domain, aa 334-417) were constructed by PCR at the positions corresponding to these domains in human/mouse ADIP ^37^. These were subcloned in-frame into pCS105-HA-RFP-ADIP and pXT7-GFP-ADIP with specific primers. Detailed cloning procedures are available upon request. All constructs have been verified by sequencing. ADIP ortholog sequences were aligned using the MEGA software, viewed using Jalview ^78^.

### Xenopus embryo culture, morpholinos, RNA synthesis and microinjections

Wild-type *Xenopus laevis* were purchased from Xenopus 1 (Michigan) and maintained in accordance with the guidelines of the Institutional Animal Care and Use Committee (IACUC) at the Icahn School of Medicine at Mount Sinai. *In vitro* fertilization and embryo culture were carried out as previously described ^59^, and embryonic staging was determined according to ^79^. Capped mRNAs were synthesized using the mMessage mMachine SP6 or T7 Transcription Kit (Thermo Fisher Scientific) and purified with the RNeasy Mini Kit (Qiagen). For microinjections, embryos were transferred to 3% Ficoll 400 (Sigma) in 0.5× Marc’s modified Ringer’s (MMR) solution (50 mM NaCl, 1 mM KCl, 1 mM CaCl_2_, 0.5 mM MgCl_2_ and 2.5 mM HEPES (pH 7.4) ^80^. Control MO (5′-GCTTCAGCTAGTGACACATGCAT-3′) ^39^, ADIP/SSX2IP MO (5’-TAACTCCTCGACTCCTTCTGGACAG-3’) ^39^, and Diversin MO (5’-GGC CAC ATC CTG CTG GCT CAT GAA T-3’) ^62^ were described previously.

To investigate protein distribution, a ventral-animal blastomere was co-injected with 100 pg of mRNAs encoding ADIP or Diversin, along with 50 pg of myrBFP. For studies on apical constriction-induced PCP, the right ventral-animal blastomere was injected with 100 pg of ADIP and 50 pg of myrBFP RNA, while the left ventral-animal blastomere was injected with 20 pg of myc-Shroom3 or Plekhg5 RNA together with 20 pg of memCherry mRNAs at the 4-cell stage. For knockdown experiments in the neural plate, a dorsal-animal blastomere was injected with 20 ng of control or ADIP morpholinos (MOs) with 50 pg of myrBFP RNA as lineage tracer. For knockdown experiments of wound closure, two ventral-animal blastomeres were injected with 40-60 ng of ADIP MOs with 100 pg of myrBFP RNA or 40 ng of Diversin MO. Embryos injected with RNA or MOs were cultured in 0.1x MMR until they reached the early gastrula or neurula stage. Each experimental group comprised a minimum of 10-20 embryos, and the experiments were repeated at least three times to ensure reproducibility.

### The wound healing and the aspiration assay

A small wound was created on the embryonic ectoderm at stage 11.5 or 14 with the dissection forceps, and wounded embryos were healed for 1-30 min in 0.7x MMR solution. The wound closure rate was monitored using a light microscope, with images captured every 5 min as previously described ^53,81^.

The *hydraulic* aspiration assay was as described by Chien et al. ^29^ with modifications. The micromanipulator holding a syringe with 0.1x MMR buffer was connected to a capillary filled with and submerged into 0.1x MMR solution. Stage 10.5-11 embryos were devitellinized and positioned with the animal hemisphere towards the capillary. Negative pressure was applied by lowering the position of the capillary to gradually draw the liquid and the embryo into the capillary, where they were subjected to suction for 25 min. Embryos exhibiting a persistent bulge post-suction were immediately fixed in MEMFA (100 mM MOPS, pH 7.4, 2 mM EGTA, 1 mM MgSO_4_, 3.7% formaldehyde) for 30 min, as described ^82^.

### ADIP antibody generation

Xenopus ADIP-specific immunogens have been produced in E. coli using glutathione-S-transferase (GST)-peptide fusions by subcloning ADIP protein fragments (amino acids 1-297, 253-384 and 295-554) into pGEX plasmids ^61^. Recombinant proteins were expressed in E. coli, purified by affinity chromatography and used to immunize Balb/c mice. Specific hybridomas were obtained after fusing splenocytes of immunized mice with the myeloma line SP-20. The hybridomas were screened by immunofluorescence on HEK-293T cells expressing full length *Xenopus* ADIP fused to RFP. Positive hybridomas were further tested by capillary immunoblotting by comparing lysates of control and RFP-ADIP-transfected HEK293T cells as previously described ^83^. Positive clones were expanded and frozen. Further testing resulted in the selection of the 5C3 antibody that recognized a single band of 65 kDa ADIP protein and was reactive with overexpressed ADIP constructs.

### Immunoprecipitation, immunoblotting and proximity labeling

Immunoprecipitations and immunoblotting have been carried out as previously described ^84^. For proximity biotinylation, four-cell embryos were injected into the animal pole with 200 pg of Flag-TurboID-ADIP and 400 pg of HA-RFP-Diversin RNAs, followed by injection of 20 nl of a 0.8 mM biotin (Millipore Sigma) solution into the blastocoel. Embryos were lysed at stage 10.5, and protein biotinylation was assessed in embryo lysates precipitated with mouse anti-HA (12A5). Immunoblotting was carried out with goat anti-biotin-HRP (Cell Signaling), anti-FLAG (M2, Sigma), mouse anti-HA (12A5), mouse anti-αTubulin (12G10, DSHB), and mouse anti-ADIP antibodies (5C3, this study). Chemiluminescence signals were measured using ChemiDoc (BioRad).

### Immunostaining and embryo imaging

For phalloidin staining, embryos were fixed in MEMFA for 1 hour at RT. Embryos were permeabilized in 0.1% Triton X-100 in PBS for 10 min and stained with Alexa Fluor 555-conjugated phalloidin (Thermo Fisher Scientific) in PBS with 1% BSA overnight at 4°C. For HA and Vangl2 staining, embryos were fixed in 2% TCA solution for 30 min at room temperature, permeabilized in 0.3% Triton X-100 in PBS for 30 minutes, and stained with mouse monoclonal anti-HA (12A5), mouse anti-E-cadherin (5D3, DSHB), and rabbit anti-Vangl2 antibody ^85^. Secondary antibodies were Alexa Fluor 488-conjugated goat anti-mouse antibody (Thermo Fisher Scientific) and Cy3-conjugated donkey anti-rabbit antibody (Jackson ImmunoResearch Laboratories).

Imaging was performed using a BC43 spinning disk confocal microscope (Fusion Ver 2 image processing software, Andor, Oxford Instruments) using 20x objective lens. Experiments were repeated at least three times, with each experimental group comprising a minimum of five embryos. Z-stack images were projected into a single image using the maximum projection function in ImageJ for subsequent analysis and quantification.

### Quantification of polarity complexes and statistical analysis

Epithelial sheet images were segmented using the Cellpose algorithm ^86^. Cell borders were identified by classifying a pixel as a border-pixel if its 8 nearest or 16 next-nearest neighbors belonged to a mask with a different cell ID. For protein aggregate analysis, pixel intensity was log-transformed, normalized, and converted into a binary mask using a 20% threshold. Protein clusters were identified using OpenCV (opencv-python 4.10.0.84, Connected components algorithm), excluding clusters smaller than 2 pixels.

To quantify *cell polarization* (see also **Fig. S1**), each protein cluster (or pixel in case of pixel-based analysis) was assigned a unit vector pointing from cell center towards its center of mass. The cell polarization vector was calculated by averaging unit vectors weighted by either cluster size or the intensity of the pixel. The *magnitude of polarity* was calculated as the length of the non-weighted average of unit vectors.

To quantify polarity orientation relative to the direction of pull, the “force vector”, defined as the shortest path from each cell center to the junction with the cell exerting the pulling force was calculated, and the angular difference between the cell’s polarization vector and the force vector was computed. Cells within the perturbed region or with incomplete boundaries were filtered out. The angular differences were binned into 12 bins and a chi-square test was used to assess the statistical significance of non-uniform angle distribution under the null hypothesis that cell polarizations orient randomly ^87^. Samples showing *p* < 0.05 were considered polarized. In ADIP-cluster analysis the null hypothesis was uniform distribution of the angles. In pixel-based analysis, the polarity quantification may be sensitive to elongated cell shape due to higher numbers of points/unit vectors. To account for this bias, a null model distribution was obtained computationally by randomly reshuffling the pixels and calculating the resulting orientation of polarity, and the corresponding rose plots were rescaled to be visually consistent with others plots. Each bin count was rescaled as 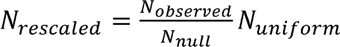, where *N_observed_* is the observed number of counts in the bin, *N_null_* the counts in the bin under the null model, and *N_uniform_* the expected counts without cell shape distortion (assumed to be uniform, *N_total_* /12 for 12 bins). For isotropic cell geometry, *N_rescaled =_ N_observed_*

*Population polarity* represents the degree of alignment of cell’s polarities across the population. The magnitude of population polarity was calculated similarly to the magnitude of cell polarity above (also Fig. S1), with the only difference of unit vectors representing the polarity orientation in individual cells. To assess whether population polarities differed significantly between samples, Welch’s t-test was applied to bootstrapped (with replacement) distributions of the population polarity.

To estimate the changes in ADIP localization over time, the aggregates overlapping with the bicellular junctions or within 1.8 µm of the tricellular junctions were counted towards bicellular, f_bi, or tricellular, f_tri, fraction, respectively. The fraction of cytoplasmic aggregates f_cyto = 1 – f_bi – f_tri.

Cell borders were imported into ImageJ as 1-pixel wide lines to measure the intensity of F-actin and E-cadherin at bicellular junctions and wound edge. Colocalization of ADIP and Diversin puncta was analyzed using the ComDet plugin in ImageJ and the scatter plots were generated using Microsoft Excel.

