## Supplemental information for "Mechanical cues organize planar cell polarity during vertebrate morphogenesis and embryonic wound repair"

**Document S1**. Figures S1–S10.

**Video S1.** ADIP polarization in response to apical constriction of neighboring cells, related to Figure 2.

Time-lapse images of ectodermal cells at stage 12 showing directional movement of RFP-ADIP (red) toward the apically constricting Shroom3-expressing cells (cyan). Images were taken every 2 mins.

**Video S2.** ADIP polarization in response to wounding, related to Figure 3**.**

Time-lapse images of gastrula ectoderm at stage 11 showing directional movement of GFP-ADIP (arrows) toward the wound (w). Asterisks mark ADIP puncta appearance near the tricellular junctions. Images were taken every 1 min.

**Supplemental Information**

**Supplemental Figures**

**Fig. S1**. **Quantification of cell polarity based on protein cluster distribution**. Related to Fig.1.

(A) Cluster-based analysis: Each protein cluster (green circle) was assigned a unit vector, **u,** pointing from cell center (black dot) towards its center of mass. (B) Pixel-based analysis: each pixel (green square) with intensity above 50% threshold of the intensity was assigned a unit vector, **u,** pointing from cell center (black dot) towards its center of mass. The cell polarization vector, **p**, was calculated by averaging unit vectors weighted by cluster size (A) or pixel-intensity (B), $w$, as $\boldsymbol{p} = \sum w_i \hat{u}_i$ . Magnitude of polarity is $m = |\sum_i^n \hat{u}_i|$To quantify polarity orientation relative to the direction of pull, the “force vector” **f**, defined as the shortest path from each cell center to the border of the cell exerting the pulling force was calculated, and the angular difference, $\alpha$, between the cell's polarization vector and the force vector was computed $\alpha = atan2((\boldsymbol{p}-\boldsymbol{f})_x, (\boldsymbol{p}-\boldsymbol{f})_y)$.

**Fig. S2. ADIP polarization is abolished by the inhibition of Myosin II activity.** Related to Fig.1.

(A) Scheme of injection. The left dorsal blastomere was injected with GFP-ADIP (100 pg), Mypt1TA (100 pg), and myr-BFP (50 pg) RNAs, and the right dorsal blastomere with GFP-ADIP (100 pg) and mem-Cherry (50 pg) RNAs. mem-Cherry and myr-BFP mark the cell boundaries.

(B) Top view of the anterior neural plate of an injected embryo at stage 16. The neural plate is highlighted by intense staining of F-actin (blue).

(C-D') ADIP localization in anterior neural ridge cells with (D-D') or without (C-C') the coexpression of Mypt1TA at the boxed region in (B). Arrows point to ADIP polarization toward the neural plate border (dashed lines). Note the lack of ADIP polarization or cell elongation in D-D'.

**Fig. S3. ADIP puncta are redistributed in response to Plekhg5-induced apical constriction** Related to Fig. 2.

(A) Stills of time-lapse images showing directional movement of RFP-ADIP (arrows) toward the apically constricting Shroom3-expressing cells marked by myrBFP. See Video S1. (B, E) Scheme of injection. The left ventral blastomere was injected with 50 pg of mem-Cherry RNA alone (B) or plus 50 pg of Plekhg5 RNA (E), and the right ventral blastomere with GFP-ADIP (100 pg) and myr-BFP (50 pg) RNAs. mem-Cherry and myr-BFP mark the cell boundaries.

(C-C'', F-F'') Distribution of GFP-ADIP (green) in cells adjacent to mem-Cherry-expressing cells (cyan) with (F-F") or without (C-C") Plekhg5 at stage 11.5.

(D, G) Rose plots showing ADIP orientation with respect to adjacent mem-Cherry-expressing cells in (C-C”) and (F-F”) respectively. 0° indicates orientation toward those cells. Data are from 3 embryos. Chi-square test indicates random distribution in (C-C”) and non-random distribution in (F-F”) respectively.

**Fig. S4. Stretching-induced ADIP polarization depends on Myosin II activity.** Related to Fig. 3.

Aspiration of gastrula embryonic ectoderm into a capillary was carried out as described in Fig. 3. (A-A”, C-C”) GFP-ADIP distribution in an unstretched (A-A”) or stretched embryo (C-C”). Note the polarization of GFP-ADIP (yellow arrows) that aligns with the stretch axis (white arrow) (C). (E-E”, G-G”) GFP-ADIP distribution in an unstretched (E-E”) or stretched embryo (G-G”) coexpressing Mypt1TA. Scale bar = 20 μm.

(B, D, F, H) Rose plots showing ADIP orientation with respect to the animal pole or aspiration axis (0°) in (A-A”), (C-C”), (E-E”), and (G-G”) respectively. Data are from 3 embryos. Chi-square test indicates random distribution in (A-A”) and (G-G”) and non-random distribution in (C-C”) and (E-E”) respectively. Red dashed lines show the expected random distribution.

**Fig. S5. Planar polarization of ADIP in multiple rows of cells surrounding the wound.** Related to Fig.4.

(A-B”’) ADIP puncta are redistributed in ectodermal cells near the wound over time. (A) Gastrula embryo with an open wound (w) in animal ectoderm. (B-B’’’) Stills of time-lapse images at the boxed area in (B) showing GFP-ADIP at 5 min (B-B’) and 41 min post-wounding (pw) (B”-B’’’), with myr-BFP marking the cell boundaries and dashed lines marking the wound edge. Note ADIP polarization toward the wound (w) at 41 min (arrows in B”-B’’’). See Video S2.

(C-C’) Localization of GFP-ADIP and myr-BFP in ectodermal cells around the wound (w) 30 min post-wounding at stage 11. Dashed line marks the wound edge, and the scale bar denotes 20 µm.

(D) Rose plot showing ADIP orientation with respect to the wound edge. 0° indicates ADIP orientation toward the wound edge. Data are from 2 embryos. Chi-square test indicates non-random distribution.

(E) Box plot showing the distribution of ADIP orientation with respect to the wound edge in each row of cells proximate to the wound edge. Red line connects median values of each group.

(F) Box plot showing the magnitude of ADIP polarization with respect to the wound edge in each row of cells proximate to the wound edge. Red line connects median values of each group.

**Fig. S6. Polarization of ADIP mutants during wound healing.** Related to Fig.4.

(A) MEGA alignment of amino acid sequences of ADIP homologs for *Homo sapiens (Q9Y2D8), Xenopus laevis (Q6NRK1), Ciona intestinalis (F7A102), Arabidopsis thaliana (Q8GW47)* and *Schizosaccharomyces pombe (Q9P6R4).* Conserved regions are highlighted. The blue line indicates the $\alpha$-Actinin-binding regions, while the magenta line represents the Afadin-binding region.

(B-D″) Localization of GFP-ADIP variants in stage 11.5 ectoderm 30 min post-wounding: full-length ADIP (B-B’), ADIP with deleted α-Actinin-binding domain (ΔAB) (C-C’) and ADIP with deleted Afadin-binding domain (ΔAFB) (D-D’). myrBFP (magenta) outlines cell boundaries. Dashed lines denote the F-actin-rich wound edge (w); arrows highlight ADIP puncta enriched at wound-proximal junctions. Scale bar, 20 µm. (E) Quantification of ADIP population polarity . Box plots represent the distributions obtained by bootstrapping, see Methods. Data are from three embryos. p-values are from Welch’s t-test.

**Figure S7. Polarization of ADIP with F-Actin and Myosin II during wound healing.** Related to Fig.4.

(A-A”) Distribution of ADIP and F-Actin in stage11 ectoderm 45 min post-wounding. Note the enrichment of ADIP (A) and F-Actin (A’) at the cell corners (arrows) facing the wound (w) edge (dashed lines).

(B-B’) Rose plots showing ADIP (B) and F-actin (B’) orientation relative to the wound edge (0°) in (A-A”). Chi-square test indicates non-random distribution compared to the null hypothesis (dashed lines).

(C-C”) Distribution of ADIP and SF9 in stage11 ectoderm 90 min post-wounding. Note the enrichment of ADIP (C) and SF9 (C’) at the cell corners (arrows) facing the wound (w) edge (dashed lines). Scale bar, 20 μm.

(D-D’) Rose plots showing ADIP (D) and SF9 (D’) orientation relative to the wound edge (0°) in (C-C”). Chi-square test indicates non-random distribution compared to the null hypothesis (dashed lines).

**Fig. S8. ADIP depletion reduces junctional F-Actin and E-Cadherin.** Related to Fig.5.

(A-B’) Epithelium of stage 14 embryos injected with indicated MOs were immunostained to visualize HA (green) and F-actin (magenta). HA-myr-BFP is the lineage tracer marking the cell boundaries.

(C) Box plots showing F-actin intensity at the control (Ctrl) junctions or wound edge at indicated time points post-wounding. Data are from three embryos with total cell numbers (n) indicated. Two-tailed Student’s t-test was performed for statistical significance.

(D-G’) Stage 14 embryos injected with indicated MOs were fixed without (D-E’) or with (F-G’) epithelial wounds (w) 5 min post-wounding and immunostained to visualize HA (cyan) and E-cadherin (green). HA-myr-BFP is the lineage tracer marking the cell boundaries, and dashed lines mark the wound edge.

(H) Box plots showing E-cadherin intensity at the control (Ctrl) junctions or wound edge. Data are from three embryos with total cell numbers (n) indicated. Mean values of each group are marked by crosses. Two-tailed Student's t-test was performed for statistical significance. n.s.: not significant.

**Fig. S9. ADIP depletion reduces junctional Vangl2.** Related to Fig.5.

(A) Stage 16 embryo unilaterally injected with ADIPMO shows neural tube closure defect (arrowhead) on the injected side (left). The anterior (A)-posterior (P) body axis is shown.

(B-C') Vangl2 immunostaining at the boxed area in (A) of embryos injected with indicated MOs. HA-myr-BFP (anti-HA staining, green) is the lineage tracer marking the cell boundaries. Note Vangl2 polarization in CoMO-injected cells (arrows in B-B') and lack of Vangl2 polarization (asterisks in C-C') in ADIPMO-injected cells.

(D) Immunoblot showing endogenous ADIP and Vangl2 in embryo lysates with indicated MOs. MAPK is the loading control.

(E-E') Localization of endogenous Vangl2 around an epithelial wound (w) 20 mins post-wounding in stage 14 embryo injected with HA-myr-BFP (green) as a membrane marker. Double immunostaining for Vangl2 and HA tag is shown. Dashed line marks the wound edge. Note the lack of Vangl2 polarization in cells around the wound.

**Fig. S10. The ankyrin repeat region is required for the colocalization of Diversin with ADIP and its polarization during wound healing.** Related to Fig.6.

(A, C) Localization of full-length RFP-Diversin (A, B) or RFP-Diversin without the ankyrin repeat domain (ΔAnk) (C, D) in the absence (A, C) or presence (B, D) of ADIP. Arrows point to puncta with Diversin and ADIP signals. (A-D) Representative stage 11 ectoderm images are shown. Cell boundaries are marked by myr-BFP (cyan). Data are from three experiments.

(E-H) Ankyrin repeats are essential for Diversin polarization. (E-E', G-G') Localization of RFP-Diversin full-length (E-E') or ΔAnk (G-G') in stage 11.5 ectoderm 30 min post-wounding. myr-BFP marks cell boundaries. Arrows point to polarized Diversin toward the wound (w) edge (dotted lines). Scale bar: 20 µm. (F, H) Rose plots showing Diversin orientation with respect to the wound edge (0°). Data are from 3 embryos. Chi-square test indicates non-random (E-E') and random (G-G') distribution.

**Supplemental videos**

**Video S1,** related to Figure 2.

**ADIP polarization in response to apical constriction of neighboring cells**.

Time-lapse images of ectodermal cells at stage 12 showing directional movement of RFP-ADIP (red) toward the apically constricting Shroom3-expressing cells (cyan). Images were taken every 2 mins.

**Video S2,** related to Figure 3**.**

**ADIP polarization in response to wounding**.

Time-lapse images of wounded gastrula ectoderm at stage 11 showing directional movement of GFP-ADIP (arrows) toward the wound (w). Asterisks mark ADIP puncta appearance near the tricellular junctions. Images were taken every 1 min.

Fig. S1

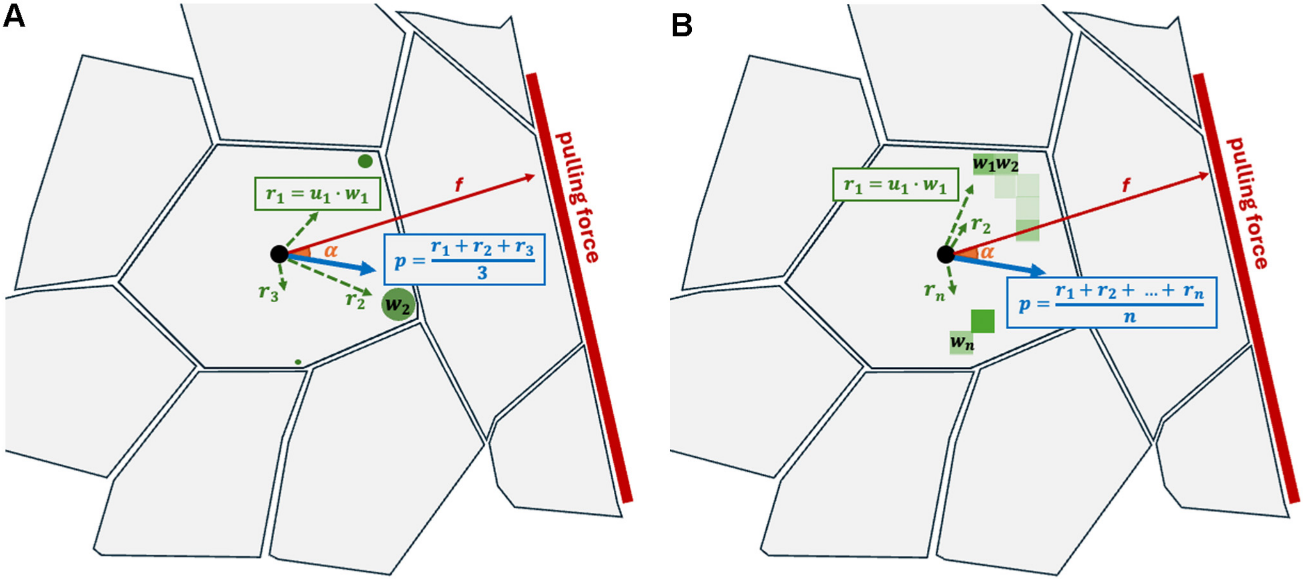

A
$r_1 = u_1 \cdot w_1$
f
α
$p = \frac{r_1 + r_2 + r_3}{3}$
$r_3$
$r_2$
$w_2$
pulling force
B
$w_1 w_2$
$r_1 = u_1 \cdot w_1$
f
$r_2$
α
$r_n$
$p = \frac{r_1 + r_2 + \ldots + r_n}{n}$
$w_n$
pulling force

### Fig. S2

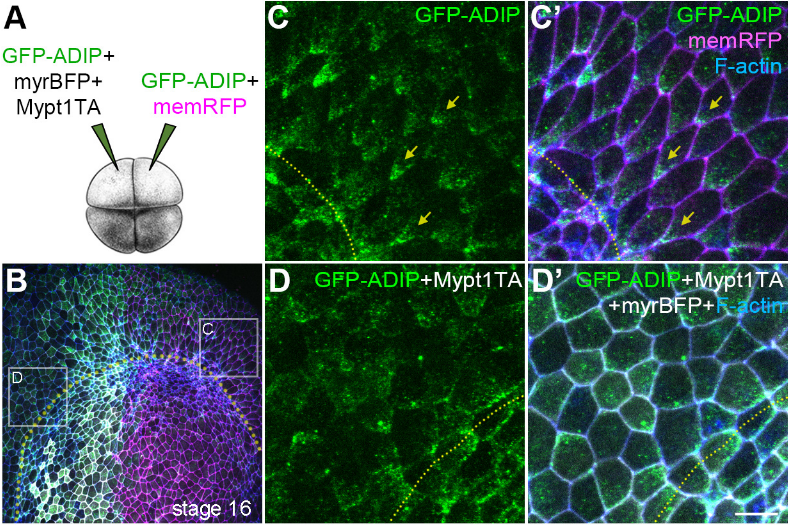
A
GFP-ADIP+
myrBFP+
Mypt1TA
GFP-ADIP+
memRFP
B
C
D
stage 16
C
GFP-ADIP
C'
GFP-ADIP
memRFP
F-actin
D
GFP-ADIP+Mypt1TA
D'
GFP-ADIP+Mypt1TA
+myrBFP+F-actin

Fig S3

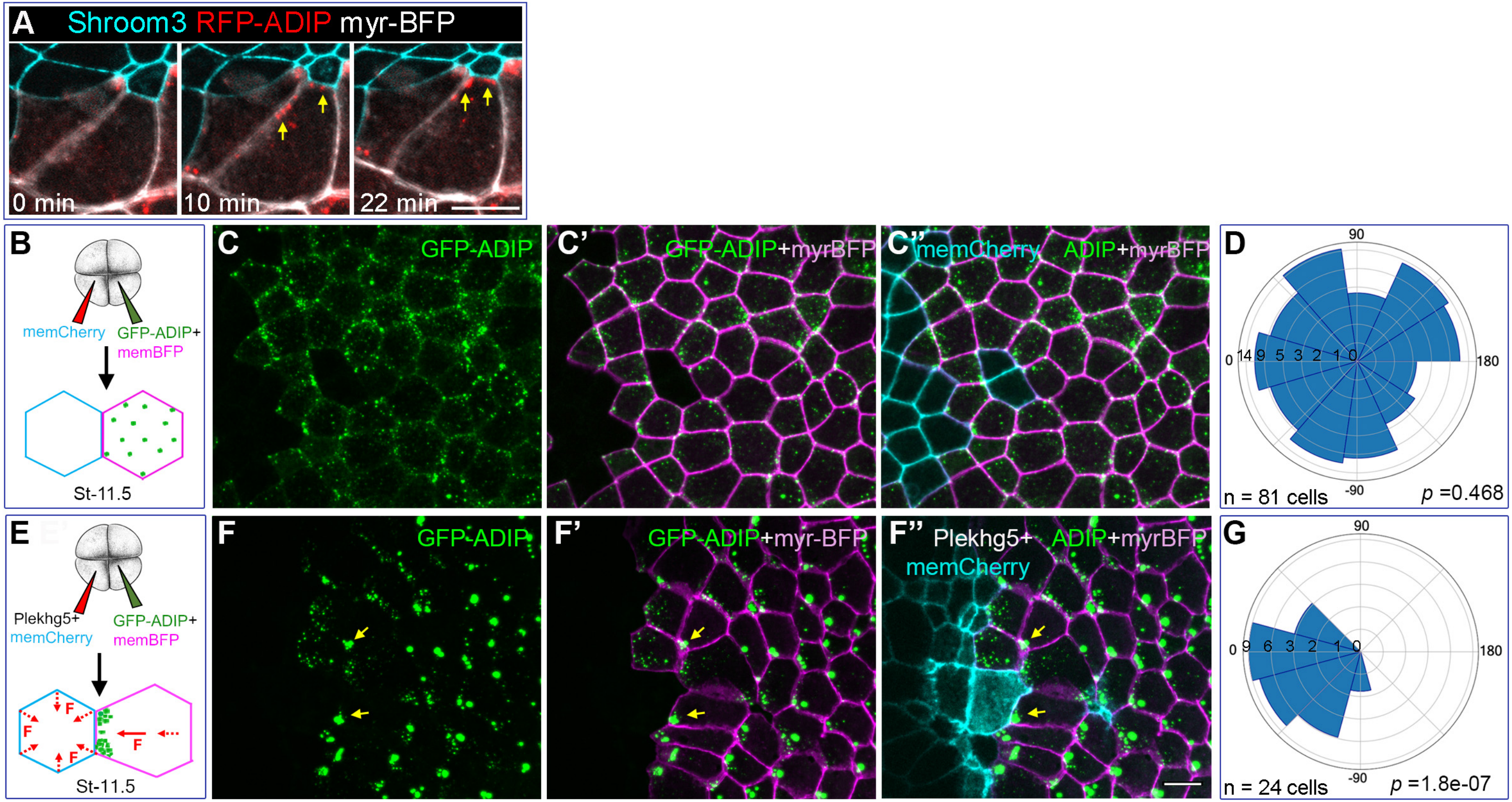
A Shroom3 RFP-ADIP myr-BFP
0 min
10 min
22 min
B
memCherry
GFP-ADIP+
memBFP
St-11.5
C GFP-ADIP
C' GFP-ADIP+myrBFP
C'' memCherry ADIP+myrBFP
D
90
0
180
-90
n = 81 cells
p =0.468
E
Plekhg5+
memCherry
GFP-ADIP+
memBFP
F
St-11.5
F GFP-ADIP
F' GFP-ADIP+myr-BFP
F'' Plekhg5+ ADIP+myrBFP
memCherry
G
90
0
180
-90
n = 24 cells
p =1.8e-07

Fig. S4

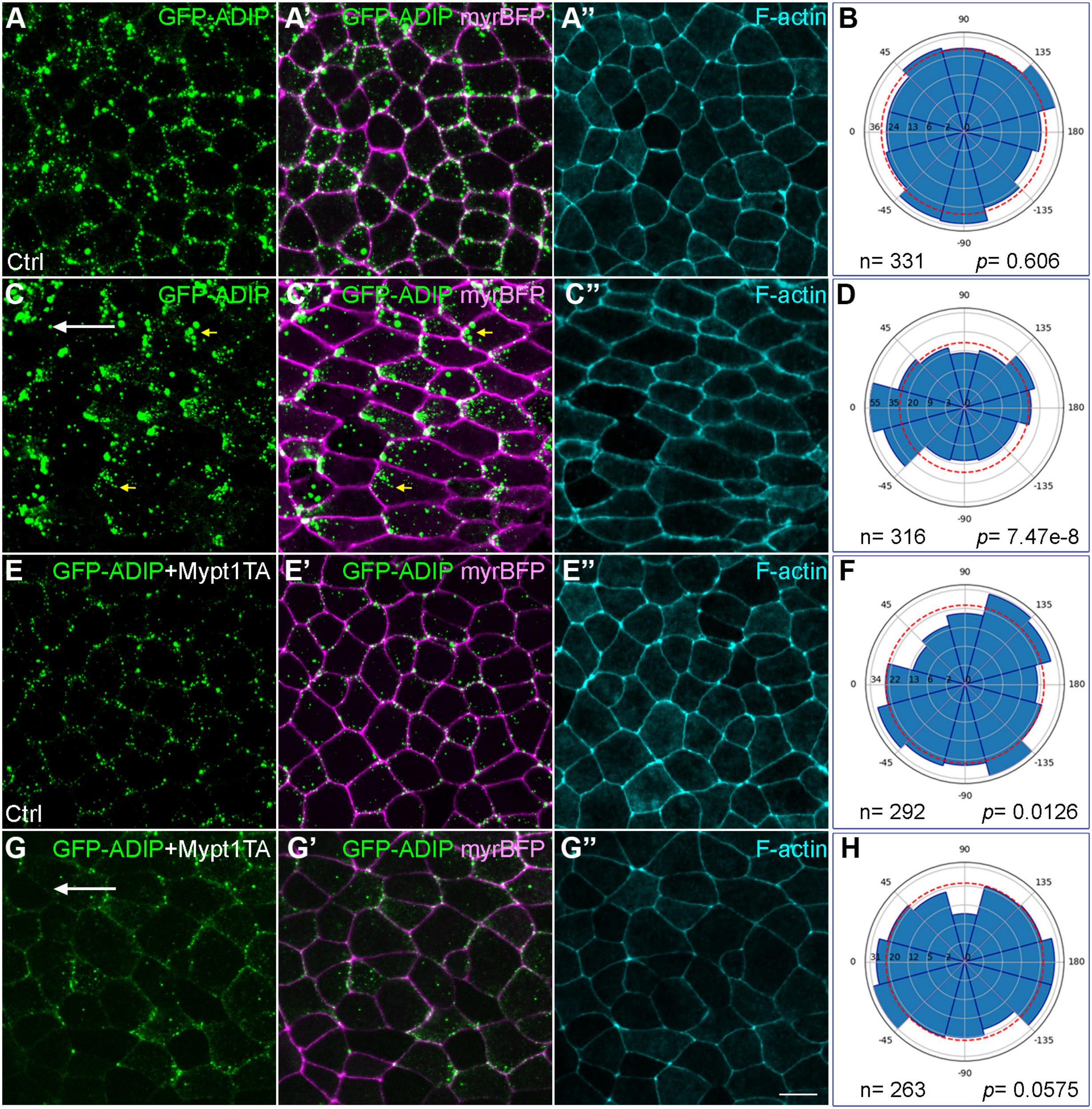
A
GFP-ADIP
Ctrl
A'
GFP-ADIP myrBFP
A''
F-actin
B
90
45
135
0
180
36
24
13
6
2
0
-45
-135
-90
n= 331
p= 0.606
C
GFP-ADIP
C'
GFP-ADIP myrBFP
C''
F-actin
D
55
35
20
9
3
n= 316
p= 7.47e-8
E
GFP-ADIP+Mypt1TA
Ctrl
E'
GFP-ADIP myrBFP
E''
F-actin
F
34
22
13
6
2
n= 292
p= 0.0126
G
GFP-ADIP+Mypt1TA
G'
GFP-ADIP myrBFP
G''
F-actin
H
31
20
12
5
2
n= 263
p= 0.0575

Fig. S5

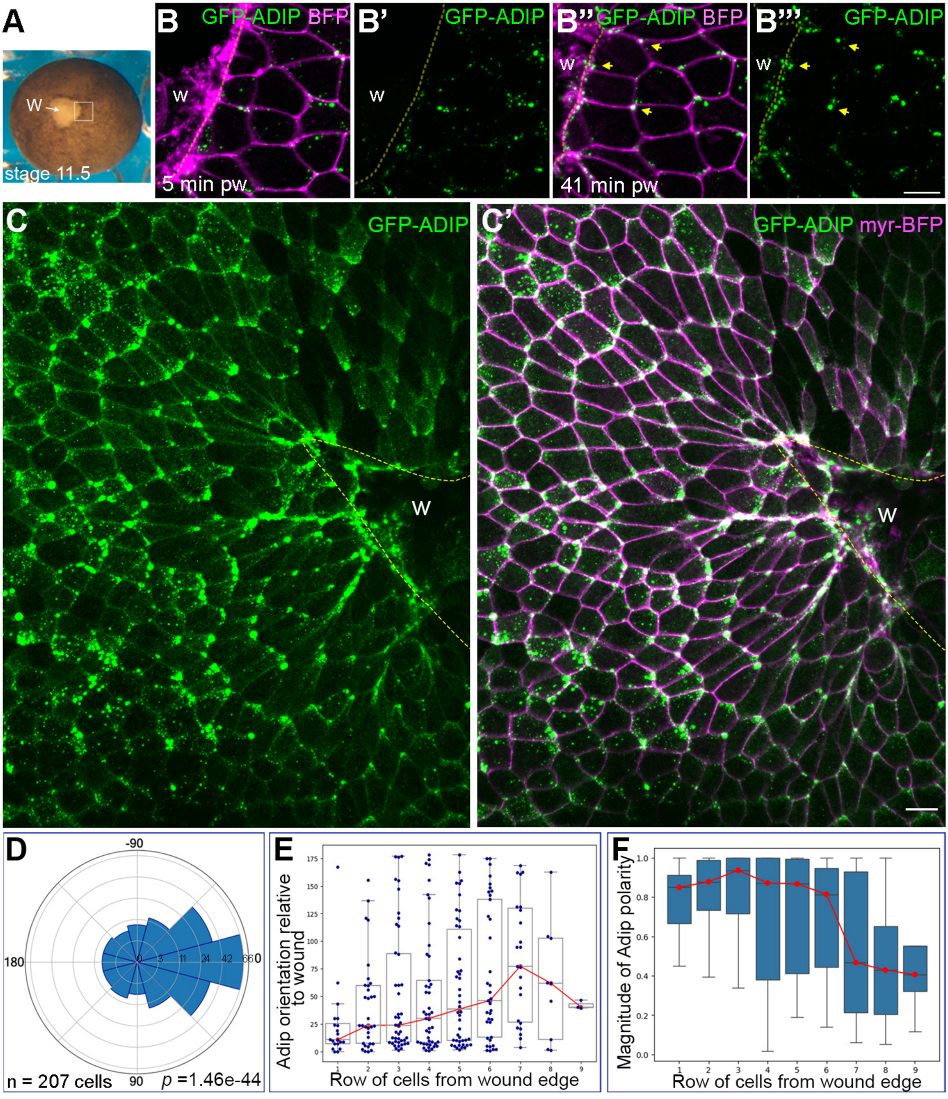
A
W
stage 11.5
B
GFP-ADIP BFP
w
5 min pw
B'
GFP-ADIP
w
B''
GFP-ADIP BFP
w
41 min pw
B'''
GFP-ADIP
w
C
GFP-ADIP
w
C'
GFP-ADIP myr-BFP
w
D
-90
180
0
90
0
3
11
24
42
66
n = 207 cells
p =1.46e-44
E
Adip orientation relative to wound
Row of cells from wound edge
F
Magnitude of Adip polarity
Row of cells from wound edge

### Fig. S6

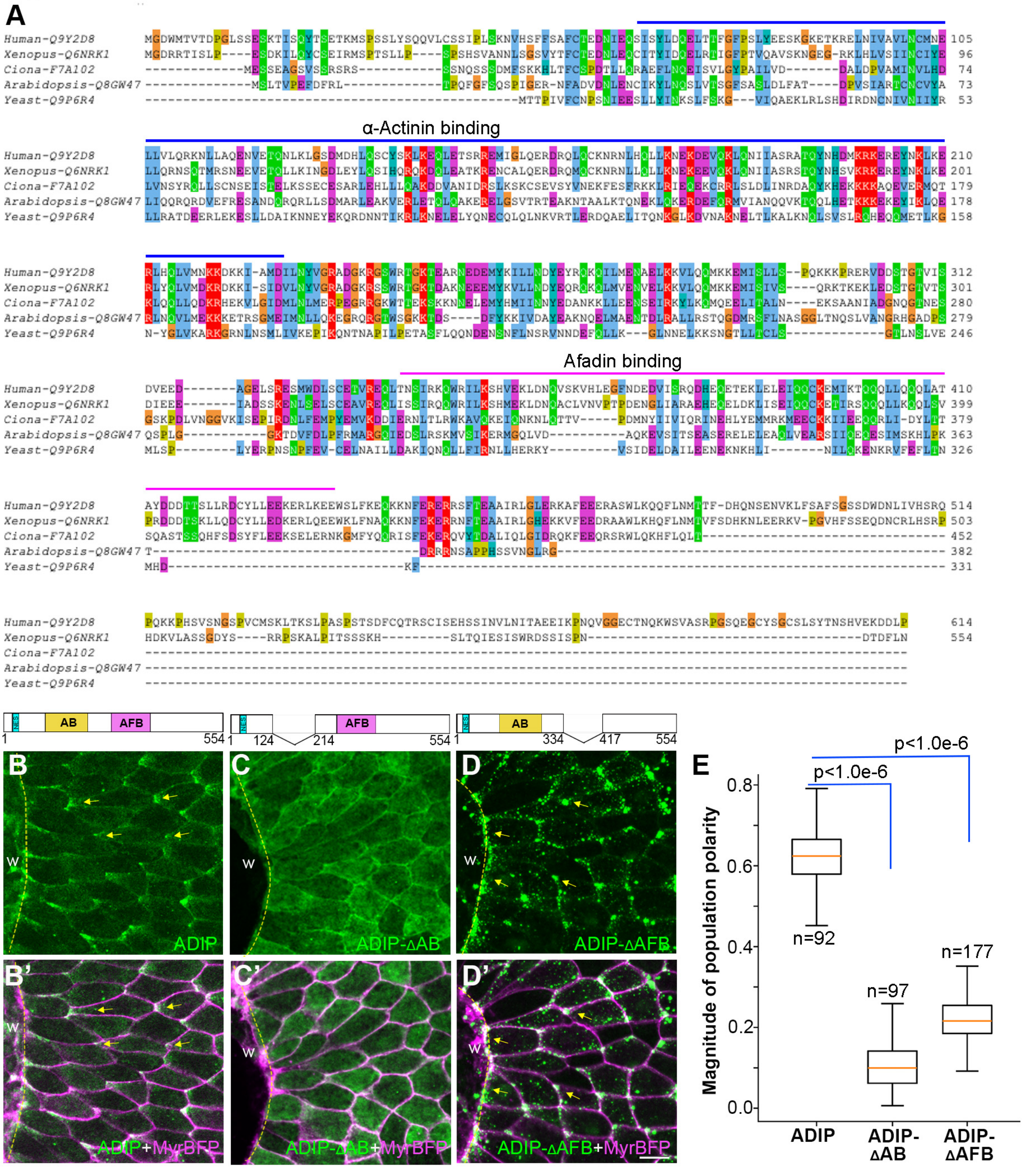
A
α-Actinin binding
Afadin binding
Human-Q9Y2D8 MGDWMTVTDPGLSSESKTISQYTSETKMSPSSLYSQQVLCSSIPLSKNVHSFFSAFCTEDNIEQSISYLDQELTTFGFPSLYEESKGKETKRELNIVAVLNCMNE 105
Xenopus-Q6NRK1 MGDRRTISLP----ESDKILQYCSEIRMSPTSLLP----SPSHSVANNLSGSVYTFCTEDNLEQCITYIDQELRTIGFPTVQAVSKNGEG-RKLHLVSIINCIYE 96
Ciona-F7A102 -------------MESSEAGSVSSRSRS-----------SSNQSSSDMFSKKHLTFCSPDTLLQRAEFLNQEISVLGYPAILVD-------DALDPVAMINVLHD 74
Arabidopsis-Q8GW47 --------------MSLTVPEFDFRL-------------TPQFGFSQSPIGER-NFADVDNLENCIKYLNQSLVTSGFSASLDLFAT----DPVSIARTCNCVYA 73
Yeast-Q9P6R4 ----------------------------------------------MTTPIVFCNPSNIEESLLYINKSLFSKG---VIQAEKLRLSHDIRDNCNIVNIIYR 53
Human-Q9Y2D8 LLVLQRKNLLAQENVETQNLKLGSDMDHLQSCYSKLKEQLETSRREMIGLQERDRQLQCKNRNLHQLLKNEKDEVQKLQNIIASRATQYNHDMKRKEREYNKLKE 210
Xenopus-Q6NRK1 LLQRNSQTMRSNEEVETQLLKINGDLEYLQSIHQRQKDQLEATKRENCALQERDRQMQCKNRNLLQLLKNEKEEVQKLQNIIASRSTQYNHSVKRKEREYNKLKE 201
Ciona-F7A102 LVNSYRQLLSCNSEISTELKSSECESARLEHLLLQAKDDVANIDRSLKSKCSEVSYVNEKFESFRKKLRIEQEKCRRLSLDLINRDAQYKHEKKKAQEVERMQT 179
Arabidopsis-Q8GW47 LIQQRQRDVEFRESANDQRQRLLSDMARLEAKVERLETQLQAKERELGSVTRTEAKNTAALKTQNEKLQKERDEFQRMVIANQQVKTQQLHETKKKEKEYIKLQE 178
Yeast-Q9P6R4 LLRATDEERLEKESLLDAIKNNEYEKQRDNNTIKRLKNELELYQNECQLQLNKVRTLERDQAELITQNKGLKDVNAKNELTLKALKNQLSVSLRQHEQMETLKG 158
Human-Q9Y2D8 RLHQLVMNKKDKKI-AMDILNYVGRADGKRGSWRTGKTEARNEDEMYKILLNDYEYRQKQILMENAELKKVLQQMKKEMISLLS--PQKKKPRERVDDSTGTVIS 312
Xenopus-Q6NRK1 RLYQLVMDKRDKKI-SIDVLNYVGRADGKRSSWRTGKTDAKNEEEMYKVLLNDYEQRQKQLMVENVELKKVLQQMKKEMISIVS----QRKTKEKLEDSTGTVTS 301
Ciona-F7A102 KLQQLLQDKRHEKVLGIDMLNLMERPEGRRGKWTTEKSKKNNELEMYHMIINNYEDANKKLLEENSEIRKYLKQMQEELITALN----EKSAANIADGNQGTNES 280
Arabidopsis-Q8GW47 RLNQVLMEKKKETRSGMEIMNLLQKEGRQRGTWSGKKTDS----DFYKKIVDAYEAKNQELMAENTDLRALLRSTQGDMRSFLNASGGLTNQSLVANGRHGADPS 279
Yeast-Q9P6R4 N-YGLVKARKGRNLNSMLIVKEPIKQNTNAPILPETASFLQQNDENSNFLNSRVNNDEFQLLK---GLNNELKKSNGTLLTCLS------------GTLNSLVE 246
Human-Q9Y2D8 DVEED-------AGELSRESMWDLSCETVREQLTNSIRKQWRILKSHVEKLDNQVSKVHLEGFNDEDVISRQDHEQETEKLELEIQQCKEMIKTQQQLLQQQLAT 410
Xenopus-Q6NRK1 DIEEE-------IADSSKENLSELSCEAVREQLISSIRQQWRILKSHMEKLDNQACLVNVPTPDENGLIARAEHEQELDKLISEIQQCKETIRSQQLLKQQLSV 399
Ciona-F7A102 GSKPDLVNGGVKISEPIRDNLFEMPYEMVKDDIERNLTLRWKAVQKEIQNKNLQTTV-----PDMNYIIVIQRINEHLYEMMRKMEECKKIIEEQQRLI-DYLTT 379
Arabidopsis-Q8GW47 QSPLG-----------GKTDVFDLPFRMARGQIEDSLRSKMVSIKERMGQLVD----------AQKEVSITSEASERELELEAQLVEARSIIQEQESIMSKHLPK 363
Yeast-Q9P6R4 MLSP--------LYERPNSNPFEV-CELNAILLDAKIQNQLLFIRNLLHERKY---------VSIDELDAILEENEKNKHLI-------NILQKENKRVFEFLTN 326
Human-Q9Y2D8 AYDDDTTSLLRDCYLLEEKERLKEEWSLFKEQKKNFERERRSFTEAAIRLGLERKAFEEERASWLKQQFLNMTTF-DHQNSENVKLFSAFSGSSDWDNLIVHSRQ 514
Xenopus-Q6NRK1 PRDDDTSKLLQDCYLLEDKERLQEEWKLFNAQKKNFEKERRNFTEAAIRLGHEKKVFEEDRAAWLKHQFLNMTVFSDHKNLEERKV-PGVHFSSEQDNCRLHSRP 503
Ciona-F7A102 SQASTSSQHFSDSYFLEEKSELERNKGMFYQQRISFEKERQVYTDALIQLGIDRQKFEEQRSRWLQKHFLQLT------------------------------- 452
Arabidopsis-Q8GW47 T------------------------------------------DRRRNSAPPHSSVNGLRG------------------------------------------ 382
Yeast-Q9P6R4 MHD-----------------------------KF-------------------------------------------------------------------- 331
Human-Q9Y2D8 PQKKPHSVSNGSPVCMSKLTKSLPASPSTSDFCQTRSCISEHSSINVLNITAEEIKPNQVGGECTNQKWSVASRPGSQEGCYSGCSLSYTNSHVEKDDLP 614
Xenopus-Q6NRK1 HDKVLASSGDYS----RRPSKALPITSSSKH---------SLTQIESISWRDSSISPN------------------------------------DTDFLN 554
Ciona-F7A102 ----------------------------------------------------------------------------------------------------
Arabidopsis-Q8GW47 ----------------------------------------------------------------------------------------------------
Yeast-Q9P6R4 ----------------------------------------------------------------------------------------------------
NES
AB
AFB
1
554
124
214
334
417
B
C
D
W
ADIP
ADIP-ΔAB
ADIP-ΔAFB
B'
C'
D'
ADIP+MyrBFP
ADIP-ΔAB+MyrBFP
ADIP-ΔAFB+MyrBFP
E
p<1.0e-6
p<1.0e-6
Magnitude of population polarity
0.8
0.6
0.4
0.2
0.0
n=92
n=97
n=177
ADIP
ADIP-ΔAB
ADIP-ΔAFB

Fig S7

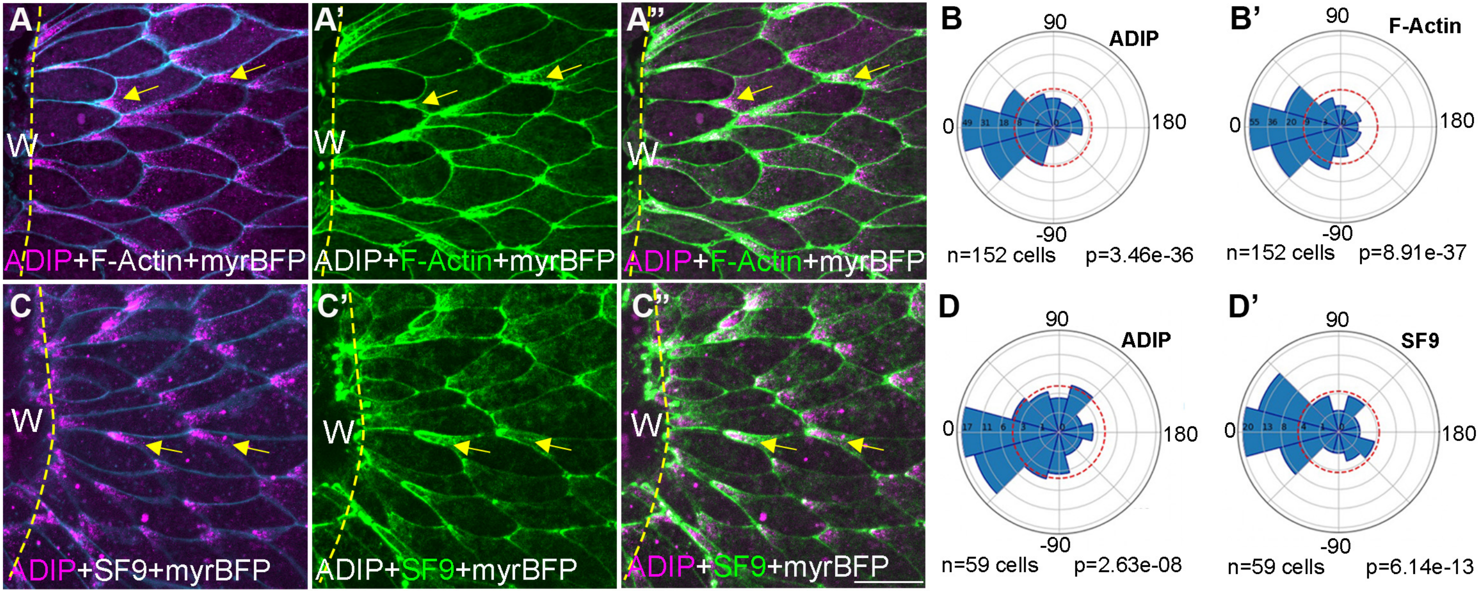

A
A'
A''
W
ADIP+F-Actin+myrBFP
B
B'
90
0
180
-90
ADIP
F-Actin
n=152 cells
p=3.46e-36
p=8.91e-37
C
C'
C''
ADIP+SF9+myrBFP
D
D'
ADIP
SF9
n=59 cells
p=2.63e-08
p=6.14e-13

Fig. S8

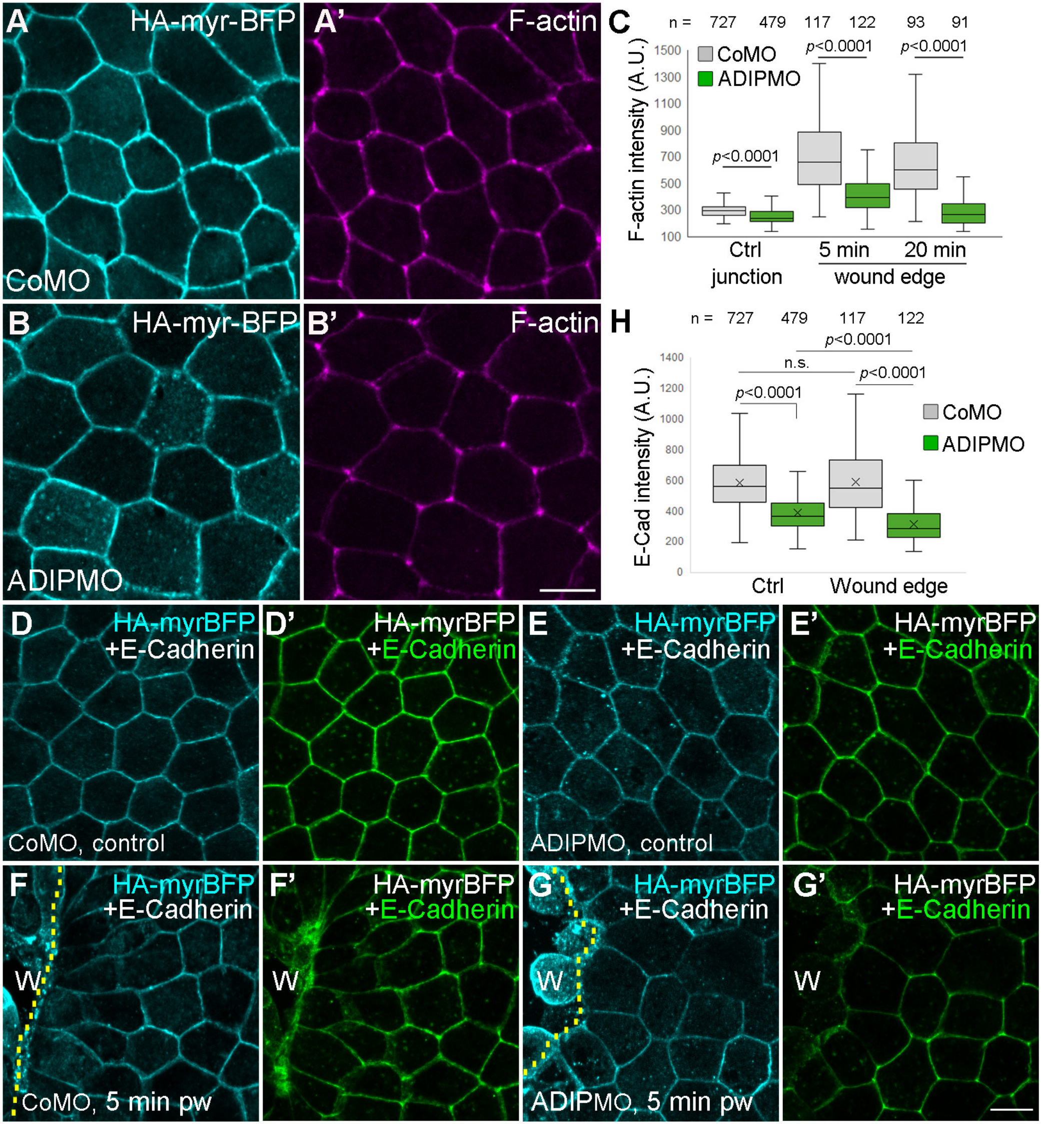
A
HA-myr-BFP
CoMO
A'
F-actin
B
HA-myr-BFP
ADIPMO
B'
F-actin
C
n = 727 479 117 122 93 91
CoMO
ADIPMO
p<0.0001
p<0.0001
p<0.0001
F-actin intensity (A.U.)
1500
1300
1100
900
700
500
300
100
Ctrl
junction
5 min
20 min
wound edge
H
n = 727 479 117 122
p<0.0001
n.s.
p<0.0001
p<0.0001
E-Cad intensity (A.U.)
1400
1200
1000
800
600
400
200
0
CoMO
ADIPMO
Ctrl
Wound edge
D
HA-myrBFP
+E-Cadherin
CoMO, control
D'
HA-myrBFP
+E-Cadherin
E
HA-myrBFP
+E-Cadherin
ADIPMO, control
E'
HA-myrBFP
+E-Cadherin
F
HA-myrBFP
+E-Cadherin
W
CoMO, 5 min pw
F'
HA-myrBFP
+E-Cadherin
W
G
HA-myrBFP
+E-Cadherin
W
ADIPMO, 5 min pw
G'
HA-myrBFP
+E-Cadherin
W

#### Fig. S9

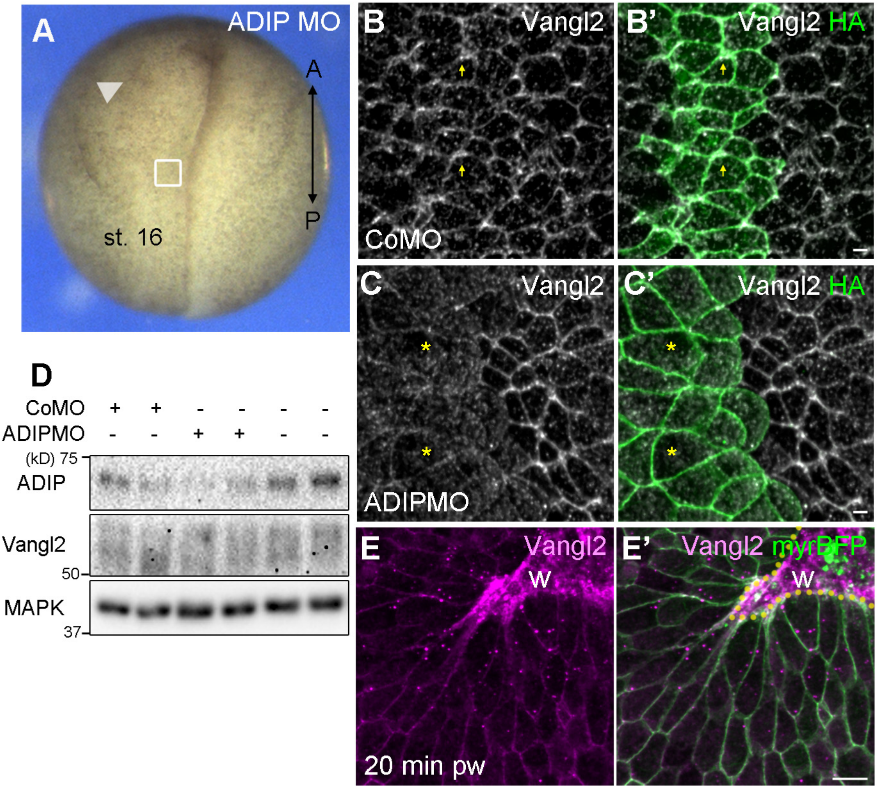

A
ADIP MO
A
P
st. 16
B
Vangl2
B'
Vangl2 HA
CoMO
C
Vangl2
C'
Vangl2 HA
ADIPMO
D
CoMO + + - - - -
ADIPMO - - + + - -
(kD) 75
ADIP
Vangl2
50
MAPK
37
E
Vangl2
W
20 min pw
E'
Vangl2 myrBFP
W

Fig. S10

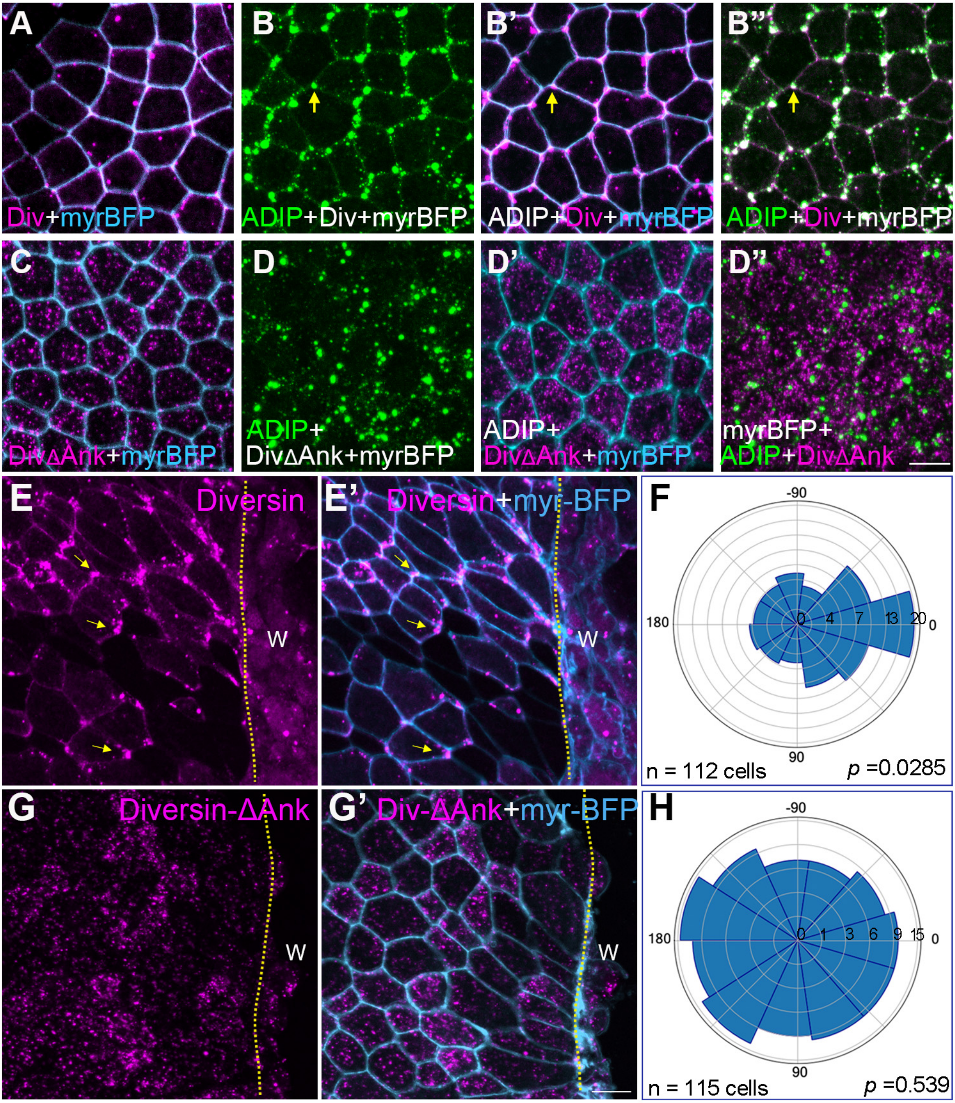
A
Div+myrBFP
B
ADIP+Div+myrBFP
B'
ADIP+Div+myrBFP
B''
ADIP+Div+myrBFP
C
DivΔAnk+myrBFP
D
ADIP+
DivΔAnk+myrBFP
D'
ADIP+
DivΔAnk+myrBFP
D''
myrBFP+
ADIP+DivΔAnk
E
Diversin
W
E'
Diversin+myr-BFP
W
F
-90
180
0
90
0
4
7
13
20
n = 112 cells
p =0.0285
G
Diversin-ΔAnk
W
G'
Div-ΔAnk+myr-BFP
W
H
-90
180
0
90
0
1
3
6
9
15
n = 115 cells
p =0.539
